# Lipodystrophy and recovery are mediated by the Wnt/lipogenesis axis during skin fibrosis

**DOI:** 10.1101/2025.09.24.678215

**Authors:** Suneeti Madhavan, Piper Moore, Rohan Varshney, Miles Montegut, Qiannan Ma, Kayla Klatt, Reshmi Parameswaran, Michael C. Rudolph, Radhika P. Atit

## Abstract

Acquired lipodystrophy in the dermal white adipose tissue (DWAT) is a salient feature of skin fibrosis, and is followed by accumulation of extracellular matrix (ECM). Lipodystrophy syndromes, often associated with metabolic co-morbidities, are estimated to affect 1 in 20,000 people. We recently showed that fibrosis-associated lipodystrophy is dependent on sustained Wnt signaling, but the mechanism is unclear. Transcriptomic profiling of mature dermal adipocytes *in vivo* reveal that Wnt activation downregulates the *de novo*-lipogenesis (DNL) axis enzymes within 48 hours. We found that protein expression of Fatty Acid Synthase (FASN), a key DNL enzyme, is dependent on sustained Wnt activation *in vitro* and *in vivo*. In human systemic sclerosis, *FASN* mRNA is significantly downregulated during two years of disease. Remarkably, pharmacological inhibition of FASN enzyme during reversal from Wnt induced fibrosis impedes recovery of DWAT lipid content as well as ECM accumulation and topography. All together, we demonstrate that acquired lipodystrophy in the context of skin fibrosis is mediated by a new role of the Wnt-DNL axis. These findings underscore the importance of this pathway in lipodystrophy and fibrosis, opening new avenues for therapeutic targets in skin fibrosis.

## 1. INTRODUCTION

Lipodystrophy, the pathological loss of adipose tissue, may be congenital or acquired, and is frequently observed alongside metabolic and autoimmune conditions such as hepatic steatosis, diabetes mellitus, Polycystic Ovarian Syndrome, Systemic sclerosis (SSc) (1). Although estimated to affect at least 1 in 20,000 people in the US, lipodystrophy often goes undiagnosed (2,3). Undetected lipodystrophy increases the risks of morbidity and mortality by disrupting homeostatic metabolism, cellular signaling, tissue elasticity, and structural integrity (4). Acquired forms of lipodystrophy, usually accompanied by metabolic abnormalities in adipocytes such as hyperglycemia, dyslipidemia and severe insulin resistance, often precedes tissue fibrosis of lung, liver, and skin in SSc (4–7). These and other fibrosis-related diseases affect approximately 1 in 4 people globally, yet no approved therapies currently exist to reverse either lipodystrophy or fibrosis (1,8). Lipodystrophy, in part, results from an imbalance of lipid uptake and lipid breakdown in adipocytes, lipofibroblasts, and related stromal cell types (4,5). Within the context of fibrosis, however, the initiating signals and cellular mechanisms that drive adipose loss remain unclear. We investigate the early signals using a novel mouse model in the skin, a tissue that is composed of matrix-rich dermis and underlying dermal white adipose tissue (DWAT).

In skin, DWAT arises from a fibro-adipoprogenitor lineage in the mouse (9,10), serving diverse physiological roles, including elasticity, wound healing, thermogenesis, hair cycle regulation, and antimicrobial defense (9,11). The loss of dermal adipocytes or their lipid content is the hallmark of multiple human skin conditions such as systemic sclerosis (SSc), androgenic alopecia, radiation- or therapy-induced cutaneous toxicity, psoriasis and skin aging (8,12–15). This DWAT lipodystrophy phenotype is also consistently recapitulated in mouse models of dermal fibrosis driven by Wnt-activated, TGFβ-activated, bleomycin and radiation exposure (16–20). In both mice and humans, lipodystrophy precedes extracellular matrix (ECM) dysfunction and contributes to fibrosis progression, resulting in thickened, stiffened, and scarred skin (15,20–23). While much of the field has focused on fibroblast activation and ECM dysregulation, the mechanisms driving DWAT depletion remain largely understudied, representing a critical opportunity to identify new molecular targets for lipodystrophy and anti-fibrotic therapies.

Acquired lipodystrophy in humans is associated with multiple signaling pathways and physiological mechanisms that impair adipose tissue function and its distribution (4,24,25). The Wnt signaling pathway stands out for its dual role as both a potent anti-adipogenic and conserved profibrotic signal across human skin, lung, and liver tissues, as well as in corresponding animal models of fibrosis [13,16,18,28–32]. Recent studies from our lab demonstrated that DWAT lipodystrophy is driven by sustained activation of canonical Wnt signaling (17,18). In this inducible and reversible genetic model, stabilized β-catenin (β-cat^istab^) is expressed in *Engrailed1* positive fibro-adipoprogenitors and their downstream dermal adipocytes, leading to lipodystrophy and ECM accumulation, both of which recover after withdrawal from doxycycline-mediated transgene activation (17,31). In parallel, ATGL-dependent lipolysis in DWAT has recently been shown to be functionally required for lipodystrophy in both bleomycin- and Wnt-induced mouse models (16,18). Importantly, in human skin SSc, lipodystrophy precedes ECM expansion, and lipid metabolic pathways are broadly dysregulated (15,16,32,33). Yet, beyond lipolysis, the specific lipid metabolic enzymes that contribute to DWAT depletion and subsequent fibrosis remain undefined.

To identify specific lipid metabolic enzymes that contributes to dermal lipodystrophy and fibrosis, we screened the transcriptome of adipocytes from the β-cat^istab^ mouse model. We discovered that *de-novo* lipogenesis (DNL), a homeostatic pathway for lipid synthesis, is downregulated during the early pathogenesis of fibrotic lipodystrophy *in vivo* and in human fibrotic diseases. Downregulation of the DNL pathway is dependent on sustained Wnt signaling activation in a cell autonomous manner in dermal adipocytes. To mechanistically connect the metabolic role of DNL to subsequent skin fibrosis, we demonstrate that activity of a principal DNL enzyme, Fatty Acid synthase (FASN), is required for the reversal of Wnt-induced DWAT lipodystrophy and ECM remodeling during fibrosis onset. Together, in the context of skin fibrosis, these findings identify the canonical WNT-DNL axis as both mechanistically important and a potential therapeutic target in dermal disease.

## 2. METHODS

### Animals

The Wnt inducible and reversible mouse model (*β*-cat^istab^) consists of a triple transgenic system which is produced by mating *Engrailed1Cre* (En1Cre); *Rosa26rTA-EGFP54* (Jax Stock 005572) and *TetO-deltaN89 β-catenin* mice as previously described (17,31). Stabilized *β*-catenin was induced in 21-day old (P21) triple transgenic mice via 6g/kg doxycycline rodent chow (Envigo-Harlan) and 2mg/mL doxycycline (Sigma) in water. After 5 days of induction, mice were euthanized with isoflurane and the dorsal dermis was harvested for intradermal adipocyte isolation, DWAT biopsy, whole skin flash freezing, paraffin or frozen sectioning. For reversal experiments, P21 mice were treated with dietary doxycycline for 21 days (until P42) and then switched to regular chow and water for 10 days, after which dorsal dermis was collected. A minimum of three mutants with litter-matched controls were analyzed for each experiment. Animals of both sexes were randomly assigned to all studies and are housed in the pathogen free animal facility maintained by the CWRU Animal Resource Centre, at the Wolstein Research Building. All the mice had access to water and feed, monitored daily by the animal husbandry technician. Mice were kept in the following environmental conditions: 12-h light/dark cycle, 20 ◦C room temperature, and 40–60 % humidity. For the bleomycin animal model, female BALB/c mice of 8-10 weeks old are purchased from the Jackson laboratory. BLM (TLC Standards catalog # B-190006) was dissolved in 0.1% sodium chloride solution at a concentration of 1mg per ml. One hundred microliters of BLM or PBS were injected subcutaneously into the back of mice daily for 14 days before sacrifice and dorsal skin collection. All animal procedures were approved by Case Western Reserve Institutional Animal Care and Use Committee (IACUC) in accordance with American Veterinary Medical Association guidelines protocol 20130156, approved 21 December 2022.

### Bulk RNA sequencing of dermal adipocytes

RNA was extracted from isolated intradermal adipocytes collected in TRIzol using the manufacturer’s protocol for high lipid, low RNA content in tissue samples. RNA-grade glycogen (Fisher Scientific, FERR0551) was used as a carrier. Total RNA quantity was measured by NanoDrop (ThermoFisher), and RNA quality was assessed by Agilent Tape Station 4200 (Agilent, Santa Clara, CA). 100 ng of total RNA was used as input to construct sequencing libraries using the Universal Plus mRNA-SEQ kit according to manufacturer protocol (NuGEN, Redwood City, CA) and sequencing data were collected with an Illumina HiSEQ4000. Sequenced reads were trimmed using trimmomatic v0.39, aligned using gsnap v2020-12-16, and filtered and sorted using SAMTools v1.9 to generate merged bam files. Raw FPKM counts were assembled, and expression levels were calculated using cufflinks v2.2.1 and the mm10 genome (GRCm38.79). A matrix of containing all samples and all genes was imported into R, filtered for genes > 1 in all samples and a per-gene row sum >=15, variance stabilization transformation was performed, and statistical differences were calculated using the DESeq2 package (v1.40.1). Data has been deposited to Gene Expression Omnibus (GEO) database under accession number GSE305616.

### In vivo FASN inhibition

FASN inhibitor TVB3664 (MCE, HY-120062) was formulated in 10% DMSO and corn oil at a concentration of 0.5 mg/mL. Triple transgenic p21 mice were put on 3 weeks of doxycycline chow diet to induce fibrosis and then moved to normal chow and water for 10 days to reverse fibrosis. During reversal, TVB3664 was administered every day via oral gavage at 5 mg/kg body weight. After 10 days of treatment, mice were sacrificed, and skin tissue was harvested as previously mentioned for further analysis.

### Immunohistochemistry

Mouse dorsal skin tissue was collected and embedded in paraffin. Paraffin sections of the skin were cut at 7μm thickness using a microtome and collected on positively charged glass microscope slides. Sections went through deparaffinization with histoclear and rehydrated with gradually decreasing graded ethanol. Then, sections were put through citrate buffer antigen retrieval (10mM sodium citrate, 1mM citric acid) for 15 minutes at 95℃. Following washes in TBS and TBS-T, slides were incubated in block buffer (1%BSA, 0.3M glycine, 0.3% Triton-X) for 1 hour at RT before incubating in rabbit-FASN primary antibody (1:1000, AbCam, ab22759) overnight. The next day, slides were incubated in either 1:1000 donkey anti-rabbit Alexa488 (for FASN-ConA) (Invitrogen, A21206) or 1:1000 donkey anti-rabbit Alexa594 (for FASN-Plin) (Invitrogen, A21207). For FASN-ConA co-stain, slides were incubated in 5μg/μL Concanavalin-A Alexa594 conjugate (Thermo, C11253) for 30 minutes before staining with 4’,6’-Diamidino-2-phenylindole (DAPI, Sigma, D9542) and mounting with Fluoroshield™ (Sigma, F6182). For FASN-Plin1 co-stain, slides were incubated in block buffer again before adding the second primary antibody for rabbit-Perilipin1 (1:1000, AbCam, ab3526) overnight. On the third day, slides were incubated in 1:1000 donkey anti-rabbit Alexa488 before staining with DAPI and mounting. Fluorescent images were taken using the Leica DMi8 inverted microscope at three separate representative regions from different sections on each slide with blue, green and red filters.

### In vivo whole skin lipidomics and triglyceride assay

Mouse skin was harvested and flash frozen immediately during dissection and whole skin lipidomics was performed using a previously described protocol (34).

### Adipocyte cell culture

Adipocytes were differentiated from isolated dermal fibroblasts from dorsal murine skin and cultured as previously described (17,35).

### In vitro FASN inhibition

FASN inhibitors TVB2640 (MCE, HY-120394) and TVB3166 (MCE, HY-112829) were formulated in DMSO. For treatment, either 10uM TVB2640 or 250uM TVB3166 was administered to differentiated adipocytes for 5 days.

### In vitro ^13^C acetate labeling, lipidomics, tracer studies and triglyceride assays

Differentiated adipocyte cultures undergoing Wnt activation and Wnt reversal were treated with 250uM sodium acetate-^13^C_2_ (Sigma, 282014) for 48 hours, with a media change every 24 hours. At end point, the cells were scraped in ice-cold acidified methanol (6:1 ratio with water and acidified with 0.1M HCl) and snap frozen. Lipidomics and tracing studies were performed in Dr. Michael Rudolph’s lab at the University of Oklahoma Health Sciences Center (OHSU) as previously described (34,36).

### Chromatin Immunoprecipitation (ChIPSeq) analysis

ChIP-seq data of TCF7L2 binding sites in human HCT116 colon cancer and human K562 acute myeloid leukemia cell lines were generated as part of the ENCODE project (GSM782123, GSM2424109) (37,38). ChIPSeq data of TCF7L2 binding sites in mouse cultured adipocytes were previously generated and are publicly available (GSE129403) (39,40). Data were obtained from Gene Expression Omnibus (GEO) database. The Integrative Genomics Viewer (IGV) was used to visualize TCF7L2 peaks.

### Transcription factor binding site (TFBS) prediction analysis

Human LEF1 motif was obtained from JASPAR (https://jaspar2020.genereg.net/) and submitted to MEMEsuite Find Individual Motif Occurrences (FIMO, https://meme-suite.org/meme/tools/fimo) and to scan *MLXIPL*, *SREBF1* and *FASN* promoter sequences (TCF7L2 bound sites from the ChIPSeq analyses) for predicted matches. CentriMo (https://meme-suite.org/meme/tools/centrimo) analyses was also performed to obtain the location and sequence of motif enrichment.

### Statistical analysis

Sample size was determined based on power analysis with 80% power. Individual data points on graphs represent the average value per mouse of 2 – 8 biological replicates (depending on measurement). Variance in standard deviation for each experiment was tested using the Barlett’s test or Brown-Forsythe test in GraphPad Prism software. Significance values when comparing two data sets were calculated by unpaired Student t-test (two-tailed) with Welch’s correction if unequal variance is detected in Prism software. Additionally, one-way analysis of variance (ANOVA) was performed on Prism to compare three or more data sets. Paired t-tests are performed where appropriate (*in-vitro* only). Significance for frequency distributions were calculated using the Kruskal Wallis test in Prism software. For graphs with individual data points, each point represents the average of one mouse. Data represent mean ± SEM. All p-values are included on the graphs and p-values less than 0.05 are considered statistically significant.

## 3. RESULTS

### 3.1 Wnt signaling activation leads to downregulation of the *de-novo* lipogenesis axis preceding dermal adipocyte lipodystrophy

Lipodystrophy of dermal adipocytes is an early and conserved feature of human and mouse fibrosis and is dependent on sustained Wnt pathway activation (15,17,18,41,42). To identify the earliest transcriptional changes induced by Wnt signaling in DWAT, we performed bulk RNA sequencing on mature dorsal skin dermal adipocytes isolated 2- and 5-days following transgene induction of βcat^istab^ to recapitulate WNT pathway activation *in vivo* (**Figure 1A**). We first confirmed effective Wnt activation responses by measuring the expression of canonical target genes, including *Axin2, Lef1,* and others that were significantly upregulated at both time points (**Supplementary Figure S1A)**. Principal Component Analysis (PCA) showed that the βcat^istab^ adipocytes express a distinct transcriptome, having many significant differentially expressed genes (DEGs) relative to control adipocytes (**Figure 1B**, **1C**). Specifically, analysis identified 475 significant DEGs at day-2 and 1319 DEGs at day-5, with 114 genes shared at both time points (*P value* ≤ 0.05) **(Supplementary Figure S1B)**. Analyses of the top 100 most significant DEGs, and the hierarchical clustering of all DEGs revealed a clear separation of DEGs responding early at 2-days and those sustained by 5-days of βcat^istab^ activation **(Supplementary Figure S1C, S1D, S1E)**, suggesting complex temporal transcriptional responses within the dermal adipocytes in our time-series.

**Figure 1:**
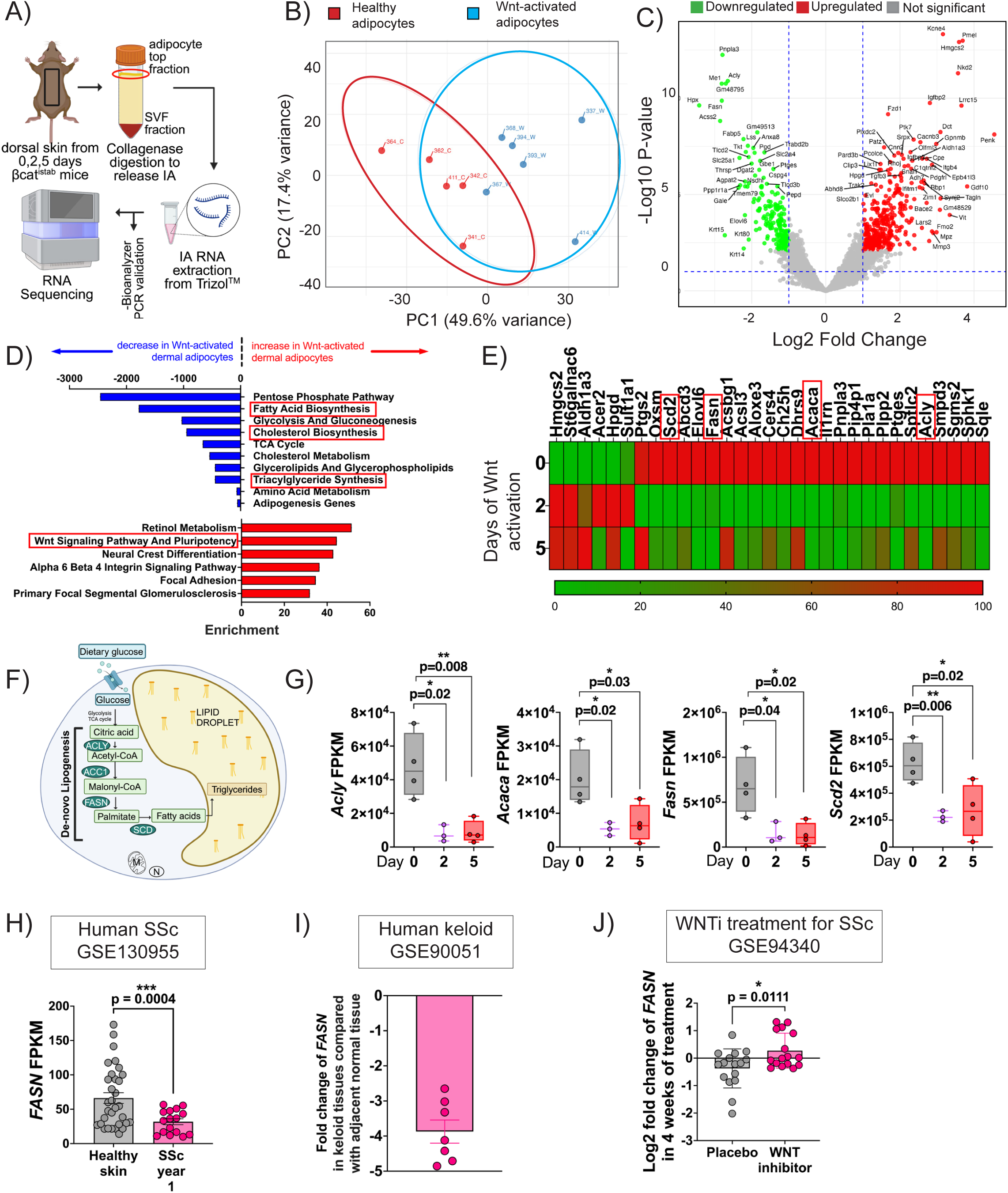
Unbiased RNA screen demonstrates that Wnt activation in adipocytes leads to downregulation of *de-novo* lipogenesis axis within 48 hours. A) Schematic of adipocyte isolation for bulk RNA sequencing. B) Principal component analysis (PCA) plot of sequenced genes from each sample grouped into healthy and Wnt-activated adipocytes. C) Volcano plot of gene expression changes (cutoff p-value <0.05) following Wnt-activation in dermal adipocytes. D) Top pathways ranked by combined odds ratio (Enrichr Wiki pathways database with p<0.05) altered in Wnt-activated dermal adipocytes compared to healthy controls. E) Heatmap of differentially expressed lipid and fatty acid metabolism genes. F) Schematic for *de-novo* lipogenesis, a process crucial for adipocytes. Excess non-lipid sugar substrate is converted to triglyceride for storage in the adipocyte lipid droplet. G) Expression (FPKM) values of enzymes in the *de-novo* lipogenesis axis indicates significant downregulation by 2 days of Wnt activation (n=4 for 0- and 5-day samples and n=3 for 2-day samples). ATP citrate lyase (*Acly*), - Acetyl Co-A carboxylase (*Acaca*), Fatty acid synthase (*Fasn*), Stearoyl-CoA desaturase (*Scd*). H) Expression of *FASN* enzyme in year 1 of human systemic sclerosis (SSc) mined from published RNA sequencing dataset GSE130955. I) *FASN* fold change in human keloid skin normalized to adjacent normal skin mined from published microarray dataset GSE90051. J) Log2 fold change of *FASN* post treatment for 28 days following placebo or topical WNT-inhibitor cream C-82 for treatment of human diffuse cutaneous SSc mined from microarray dataset GSE94340. P-values were calculated with unpaired, two-tailed t-test with Welch’s correction when required. *, **, ***, **** is p-value <0.05, 0.01, 0.001 and 0.0001 respectively. A p-value < 0.05 is considered significant.

Pathway analysis using Enrichr identified the gene ontology (GO) terms lipid biosynthesis, including fatty acid, cholesterol, and triglyceride synthesis were populated by the significantly downregulated genes following βcat^istab^ activation in dermal adipocytes **(Figure 1D)**. Consistent with effective activation (Supp Figure 1A), Enrichr identified the “Wnt Signaling Pathway and Pluripotency” GO term that was populated by upregulated DEGs (Figure 1D). Moreover, Gene Ontology analysis of the DEGs using DAVID (https://david.ncifcrf.gov/) similarly identified “Fatty Acid Biosynthetic Process” and “Lipid Metabolic Process” as the most enriched GO terms at 2-days of βcat^istab^ activation in adipocytes **(Supplementary Figure S1F)**. Interestingly, the GO terms “extracellular region” and “ECM binding” were populated with upregulated genes at 2-days of βcat^istab^ activation, with a tenfold increase in the enrichment score after 5-days **(Supplementary Figure S1F, S1G)**. These early molecular changes in lipid metabolism and subsequent robust increase in ECM signature support the kinetics of fibrosis progression observed in animal models and human skin fibrosis (17,30,42,43).

Next, we investigated the 31 DEGs enriched in the “Fatty Acid Biosynthetic Process” and “Lipid Metabolic Process” pathways following 2-days of βcat^istab^ activation in dermal adipocytes. We identified downregulation of the core genes encoding the enzymes required for *de-novo* fatty acid synthesis: ATP citrate lyase (*Acly*), Acetyl-CoA carboxylase 1 (*Acaca*), and Fatty Acid Synthase (*Fasn*) (**Figure 1E**). In addition, stearoyl-CoA desaturase 2 (*Scd2*), which catalyzes desaturation of saturated fatty acyl-CoAs, was also suppressed. Together, *Acly*, *Acaca*, *Fasn*, and *Scd2* form a component of the DNL pathway that converts carbohydrate-derived carbon into palmitate (16:0), which is then desaturated into palmitoleate (16:1) (**Figure 1F**). Significant suppression of all four genes was observed at 2-days of βcat^istab^ activation, with sustained downregulation through day-5 in dermal adipocytes (**Figure 1G and S1C**). Importantly, the key identity markers, including Peroxisome Proliferator activated receptor gamma (*Pparg*), CCAAT enhancer-binding protein alpha (*Cebpa*), Perilipin 1 (*Plin1*) and Adiponectin (*AdipoQ*), were unchanged at both time points, indicating that early Wnt activation suppressed lipid biosynthesis without altering dermal adipocyte identity (**Supplementary Figure S2A**). Surprisingly, major transcriptional regulators of the DNL program, Sterol regulatory element-binding transcription factor 1 (*Srebf1*) and MLX-interacting protein-like, encoding ChREBP/ Carbohydrate-responsive element-binding protein *(Mlxipl*), were also unchanged (**Supplementary Figure S2B**). *Mlxipl* expression was modestly reduced but did not approach statistical significance by 2-days of βcat^istab^ activation.

To assess the specificity of Wnt-mediated DNL suppression, we assessed other lipid-handling pathways. Genes involved in glucose transport solute carrier family 2 member 4; encodes GLUT4/Glucose transporter 4 (*Slc2a4*), transport of mitochondrial citrate Solute Carrier Family 25 Member 1, encoding the CIC/Mitochondrial citrate carrier (*Slc25a1*), elongation of DNL fatty acids ELOVL fatty acid elongases 5 and 6 (*Elovl5* and *Elovl6*, and triglyceride synthesis Lipin (*Lpin*) were all significantly downregulated **(Supplementary Figure S2C)**. In contrast, genes associated with fatty acid uptake and lipolysis was largely unchanged, with the exception of Caveolin 1 (*Cav1)*, Caveolin 2 (*Cav2*), and Adenyl cyclase (*Adcy1*) (**Supplementary Figures S2D and S2E**). Although prior studies have implicated fibrosis-associated metabolic reprogramming via the Warburg effect, a shift between glycolysis and fatty acid oxidation (20), no significant differences for genes in these pathways were observed (**Supplementary Figure S2F, S2G**). Together, these findings indicate that selective transcriptional suppression of the DNL axis is an early and specific molecular signature of Wnt-driven dermal dysfunction during skin fibrotic progression.

To evaluate whether DNL suppression also occurs in human fibrotic skin, we performed a meta-analysis of publicly available transcriptomic data. In the Prospective Registry of Early Systemic Sclerosis (PRESS) cohort, which includes skin biopsies from 48 patients with early, diffuse SSc and mean disease duration for ∼1.3 years (GSE 130955) (44), we found that expression of *FASN*, a rate-limiting DNL enzyme, was significantly reduced, while the other DNL pathway genes remained unchanged (**Figure 1H, Supplementary Figure S3A**). Similarly, in human keloid scar tissue, another fibrotic dermal condition, *FASN* and additional DNL axis genes were downregulated relative to healthy skin controls (GSE90051) (45) (**Figure 1I, Supplementary Figure S3B**). Finally, in a short-duration clinical trial of C-82, a topical small molecule inhibitor of Wnt/β-catenin, gene expression analysis of biopsies from SSc patients revealed partial restoration of the adipocyte gene expression signature, including significant upregulation of FASN and other lipid synthesis genes (GSE94340) (46) (**Figure 1J, Supplementary Figure S3C)**. Together, these findings demonstrate that reduction in FASN expression is an early and conserved molecular event in human fibrotic skin, observed in both SSc and keloid scarring, and that FASN expression is at least partially recoverable following Wnt pathway inhibition in human dermal fibrosis.

### 3.2 Wnt-induced downregulation of *de-novo* lipogenesis is cell autonomous to mature dermal adipocytes

To determine whether canonical Wnt signaling suppresses the DNL pathway in a cell-autonomous manner, we activated the Wnt pathway in mature mouse dermal adipocytes following differentiation of SVF in vitro, as previously described (17) (**Figure 2A**). Treatment with either CHIR99021 (a GSK-3 inhibitor; 7 μM) or BML284 (a β-catenin stabilizer; 20 μM) for 5-days resulted in robust upregulation of the Wnt target gene *Axin2*, without affecting *Plin1* mRNA levels, confirming pathway activation without loss of adipocyte identity **(Supplementary Figure S4A).** Both pharmacological treatments resulted in a significant reduction in adipocyte lipid accumulation, as measured by Oil-Red-O staining (**Figure 2B**). Consistent with our *in vivo* results, CHIR99021 treatment led to a significant decrease in *Slc2a4* (GLUT4), *Acly*, and *Fasn* transcript levels, as well as reduced FASN protein levels (**Figure 2C**, **2D; Supplementary Figure S4B**). While *Srebf1* and *Mlxipl* were not significantly altered *in vivo*, the downward trend of *Mlxipl* observed in the RNA seq screen prompted further validation of DNL transcription factors *in vitro* **(Supplementary Figure S2B**). We found that the transcriptional regulator *Mlxipl* (ChREBP) was dramatically reduced after 5 days of CHIR treatment in mature adipocytes while *Srebf1* expression was comparable to controls (**Figure 2E**). Morphometric analyses indicated significant reductions in the number, area, perimeter, and circularity of lipid droplets in CHIR-treated adipocytes compared to controls (**Figure 2F**, **2G, Supplementary Figure S4C, S4D**). These results demonstrate that pharmacological activation of Wnt signaling is sufficient to suppress DNL gene expression and diminish lipid droplet formation in an adipocyte-autonomous manner, recapitulating the transcriptional and morphological phenotype observed *in vivo*.

**Figure 2:**
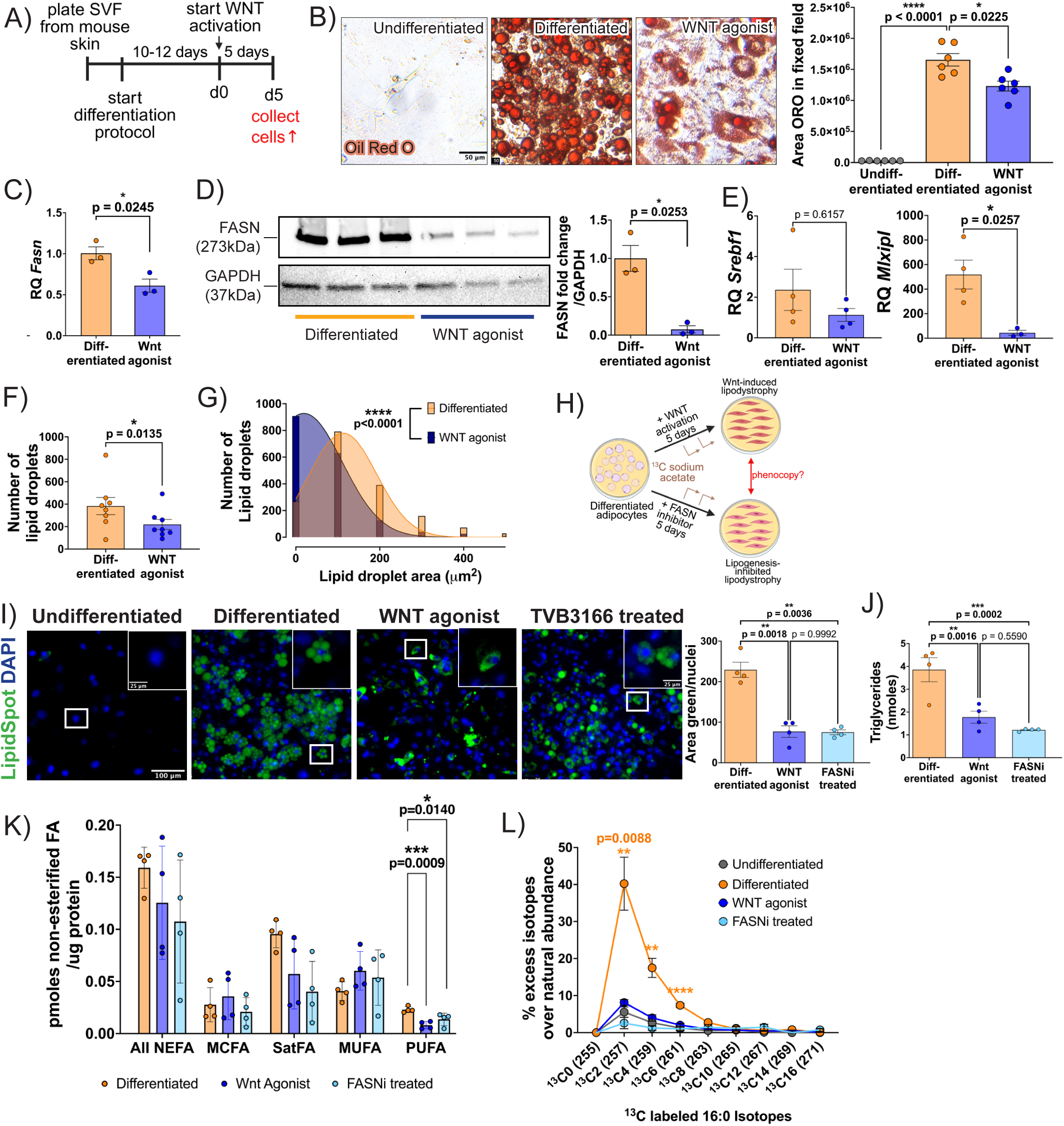
Cultured adipocytes undergo Wnt-induced downregulation of *de-novo* lipogenesis. A) Schematic of differentiated adipocyte cell culture model. B) Oil red O staining of undifferentiated, differentiated and 5-day 7µM WNT agonist CHIR99021 treated cells (scale bar = 50um) and respective image quantification of adipocyte lipid droplets using CellProfiler (n=6). C) Relative quantity of *de-novo* lipogenesis enzymes mRNA *Slc2a4 (*GLUT4), *Acly* and *Fasn* normalized to *Hprt* housekeeping gene in differentiated and WNT agonist CHIR99021 treated adipocytes (n=3). D) Band intensity and corresponding quantification of FASN protein (273 kDa) normalized to GAPDH i(n=3). E) Relative quantity of *de-novo* lipogenesis transcription factor *Mlxipl* mRNA normalized to *Hprt* (n=3-4). F) Number of lipid droplets per fixed field (n=8) quantified using CellPose. G) Histogram of individual lipid droplet area (in microns^2^) (n= 1724 randomly selected droplets). H) Schematic showing experimental design of WNT agonism, FASN inhibition and ^13^C acetate pulse (DNL assay) in the differentiated adipocyte cell culture model. I) Representative undifferentiated, differentiated, 5-day WNT agonist CHIR99021 treated and 5-day 250nM TVB3664 (FASN inhibitor) treated cultures co-stained with LipidSpot (lipid droplet, green) and DAPI (nucleus, blue) with inset (white square) showing high magnification single cell morphology (scale bar=100μm, for inset = 25μm) with corresponding graph quantifying the area of LipidSpot dye normalized to nuclei counts (n=8). J) Quantification of moles of neutral triglyceride (TAG) from cultures (n=4). K) Lipidomics analysis of WNT agonist BML-284 treated and FASN_i_ treated adipocytes showing non-esterified fatty acids (NEFA) grouped by total NEFA, medium chain FA (MCFA), mono-unsaturated FA (MUFA) and poly-unsaturated FA (PUFA) normalized to protein quantity (n=4). L) ^13^C-isotope incorporation in 16:0 (palmitate) species (n=4). For two groups, unpaired two-tailed Students t-test and for three or more groups, one-way ANOVA with appropriate posthoc test was employed. ^For the frequency distribution, non-^ parametric Kruskal-Wallis test was applied. *, **, ***, **** is p-value <0.05, 0.01, 0.001 and 0.0001 respectively. A p-value < 0.05 is considered significant.

Fatty Acid Synthase (FASN) enzyme catalyzes the seven reactions that convert carbohydrate-derived carbon in the form of acetyl-CoA and malonyl-CoA substrates into the saturated fatty acid palmitate (36,47). To determine whether loss of FASN function is sufficient to phenocopy Wnt-mediated lipodystrophy, we treated mature dermal adipocytes *in vitro* with the FASN inhibitor (FASN_i_) TVB-2640 (10μM) for 5 days (**Figure 2H)**. FASN inhibition significantly reduced neutral lipid accumulation, as assessed by LipidSpot staining, and decreased triglyceride content, lipid droplet area and perimeter, closely resembling the phenotype of Wnt-activated adipocytes treated with BML284 (**Figures 2I**, **2J; Supplementary Figures S4E–G)**. To directly assess whether DNL was functionally impaired following Wnt activation, we performed a stable isotope tracing assay using [^13^C]-acetate to quantify incorporation of labeled carbon into palmitate, as we previously described (48). Mature dermal adipocytes were incubated with BML284 or FASN_i_ for 5 days and pulsed with [^13^C]-acetate for the last 48 hours **(Figure 2H)**. While both Wnt-activated and FASN-inhibited adipocytes exhibited comparable levels of total and non-esterified fatty acids relative to controls, there was a significant reduction in non-esterified palmitate (16:0), non-esterified polyunsaturated fatty acids (PUFAs) and the saturation index (ratio of saturated to unsaturated fatty acids indicating SCD enzyme activity) (**Figure 2K, Supplementary Figure S4H, S4I, Supplementary table 1)**. Furthermore, both Wnt-activated and FASN inhibited adipocytes incorporated significantly less ^13^C tracer into the M+2, M+4, and M+6 palmitate ions. All together, these data indicate impaired DNL function and suggests that activated WNT functions upstream of FASN activity (**Figure 2L)**. Together, these results demonstrate that FASN is functionally required for lipid accumulation in dermal adipocytes, and its inhibition is sufficient to phenocopy the Wnt activation mediated dermal adipocyte lipodystrophy.

### 3.3 Wnt induced downregulation of *de-novo* lipogenesis recovers in Wnt-reversed mouse skin

Given that *FASN* is suppressed early in human SSc and keloids tissues and recovers upon Wnt inhibition in human SSc (**Figure 1H-J**), we tested whether DNL axis gene and protein expression was lost during βcat^istab^ activation and restored following reversal of βcat^istab^ activation *in vivo*. After 5-days of β-cat^istab^ induction in *vivo*, the DWAT thickness and the individual PLIN+ adipocyte area and perimeter were significantly reduced relative to controls (**Figure 3A-C Supplementary Figure S5A**), indicating that early and selective lipodystrophy ensues following activation of the β-cat^istab^ transgene. We used DWAT biopsies from intact skin and found *Fasn* and *Acaca* mRNA expression was significantly decreased, while *Plin1* was unchanged, independently confirming our isolated adipocyte bulk RNASeq data (**Figure 3D, 3E**, **Supplementary Figure S5B).** Consistent with our *in vitro* studies, we found *Srebf1* expression was comparable to controls, while *Mlxipl* was significantly downregulated in 5-day β-cat^istab^ DWAT biopsies **(Figure 3E)**.

**Figure 3:**
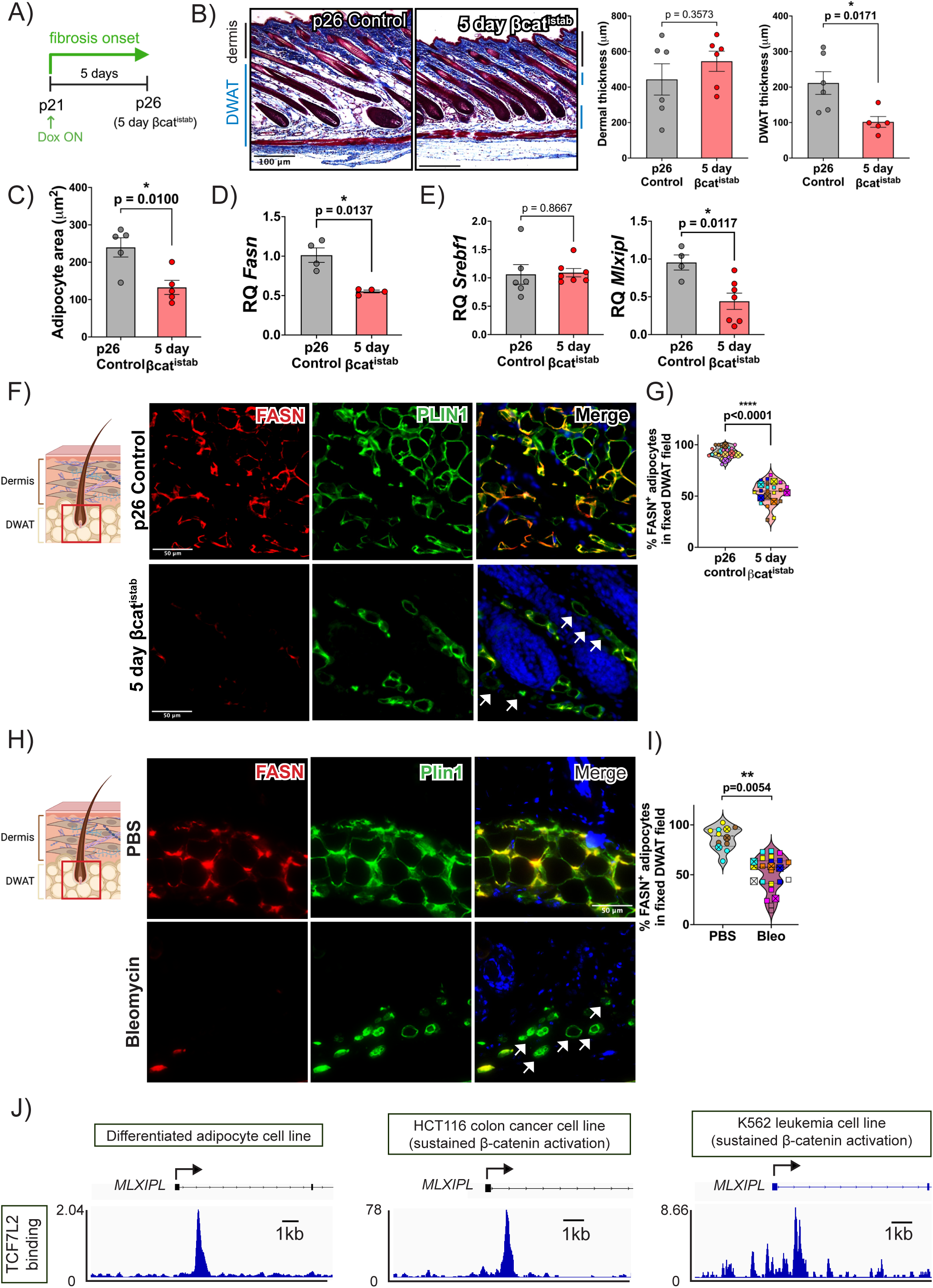
FASN is downregulated in genetic and chemical animal models of fibrosis. A) Schematic representing design of 5 days βcat^istab^ experiment. B) Histology of 5-day βcat^istab^ mouse skin with age matched control and corresponding quantification of dermal and DWAT thickness (n=6). C) Quantification of average individual adipocyte area (microns^2^) in p26 control and 5-day βcat^istab^ (n=5) D) Relative quantification of *Fasn* mRNA from DWAT biopsy tissue of both groups (n=4) normalized to *ActB*. E) Relative quantification of DNL transcription factors *Srebf1* and *Mlxipl* from DWAT tissue biopsy of both groups normalized to *ActB* (n=6-7). F) Immunofluorescence for FASN (red) and Perilipin1 (green) in Control and 5d βcat^istab^ DWAT. White arrows indicate adipocytes without FASN. G) Quantification of percent adipocytes with FASN colocalization (FASN^+^) compared to total adipocytes counted in a fixed field (n=6-7). Each large symbol is the average of three representative 40X fields (small symbol) in one animal. H) Immunofluorescence for FASN (red) and Perilipin1 (green) in PBS-treated and 14-day bleomycin (bleo) treated DWAT. White arrows indicate adipocytes without FASN. I) Respective quantification of FASN^+^ adipocytes (n=3-6). J) Integrative Genomics Viewer (IGV) view of TCF7L2 binding occupancy in regions ± 3kb from transcription start sites (arrows) of *Mlxipl* in differentiated adipocytes, HCT116 and K562 cell lines. P-values were obtained using unpaired students t-test. *, **, ***, **** is p-value <0.05, 0.01, 0.001 and 0.0001 respectively. A p-value < 0.05 is considered significant.

To validate that Wnt signaling regulates FASN abundance at the protein level *in vivo*, we assessed FASN protein in PLIN1+ dermal adipocytes within the DWAT in the β-cat^istab^ genetic model and the bleomycin (Bleo)-induced skin fibrosis model **(Supplementary Figure S5C)**. In control mice, nearly all PLIN1^+^ adipocytes expressed FASN protein. In contrast, nearly half of PLIN1^+^ dermal adipocytes lacked detectable FASN protein expression in 5-day βcat^istab^ and 14-day bleo-treated DWAT (**Figure 3F-I**). Relative fluorescence intensity of FASN in bleo-treated DWAT was also dramatically reduced compared to controls **(Supplementary Figure S5D)**.

To investigate the mechanism of β-catenin induced downregulation of DNL, we revisited published β-catenin co-factor TCF7L2 ChiP-seq studies on human colon cancer cell line HCT116 and human myeloid leukemia cell line K562 which are known to have elevated and sustained Wnt signaling (GSM782123, GSM2424109) (37,38) and human differentiated adipocyte cell line (GSM3712534) (39,40). We found a robust peak within 3kb of the transcriptional start site (TSS) of HCT116 and K562 cells that was also present in adipocytes near exon 1 of *MLXIPL* and *FASN* (**Figure 3J, Supplementary Figure S6A).** Interestingly, the other key regulator of DNL, *SREBF1* had weak or no TCF7L2 peaks within 3kb of the TSS in these cell lines **(Supplementary Figure S6A).** Since TCF7L2 and LEF1 are co-factors for Wnt/β-catenin regulation, we probed for predicted LEF1 transcription factor binding sites on the TCF7L2 bound promoter sequence of *MLXIPL*, *SREBF1* and *FASN* (49,50). We found one partially matched LEF1 motif on TCF7L2 regions of all three genes **(Supplementary Figure S6B)**. The identification of a robust TCF7L2 binding site and predicted LEF1 binding site near the *MLXIPL* gene in three different cell types suggest that ChREBP (*MLXIPL*) is a potential direct transcriptional target of Wnt/β-catenin signaling, offering a mechanistic link between Wnt activation and DNL suppression in dermal adipocytes.

We previously showed that withdrawal from Wnt activation restored lipid laden adipocytes in DWAT, and resolved the fibrotic ECM three weeks later (17). Next, we tested if downregulation of FASN requires sustained Wnt activation *in vivo*. After 10 days of reversal from β-cat^istab^ transgene activation, we observed the re-emergence and recovery of the DWAT layer thickness, individual adipocyte area, perimeter, and circularity (**Figure 4A-E**). After 10-days of βcat^istab^ reversal, the percent of DWAT adipocytes expressing FASN protein expression in PLIN1^+^ adipocytes tended to be lower but was comparable to controls, suggesting that an emergence from lipodystrophy requires withdrawal from Wnt activation **(Figure 4F)**. These results suggest that FASN suppression is a defining and reversible feature of Wnt-driven fibrotic remodeling in skin. Together, these findings demonstrate that FASN protein expression is dynamically regulated by elevated Wnt signaling in *vivo*, and that recovery from dermal lipodystrophy is accompanied by restoration of FASN expression in mature adipocytes.

**Figure 4:**
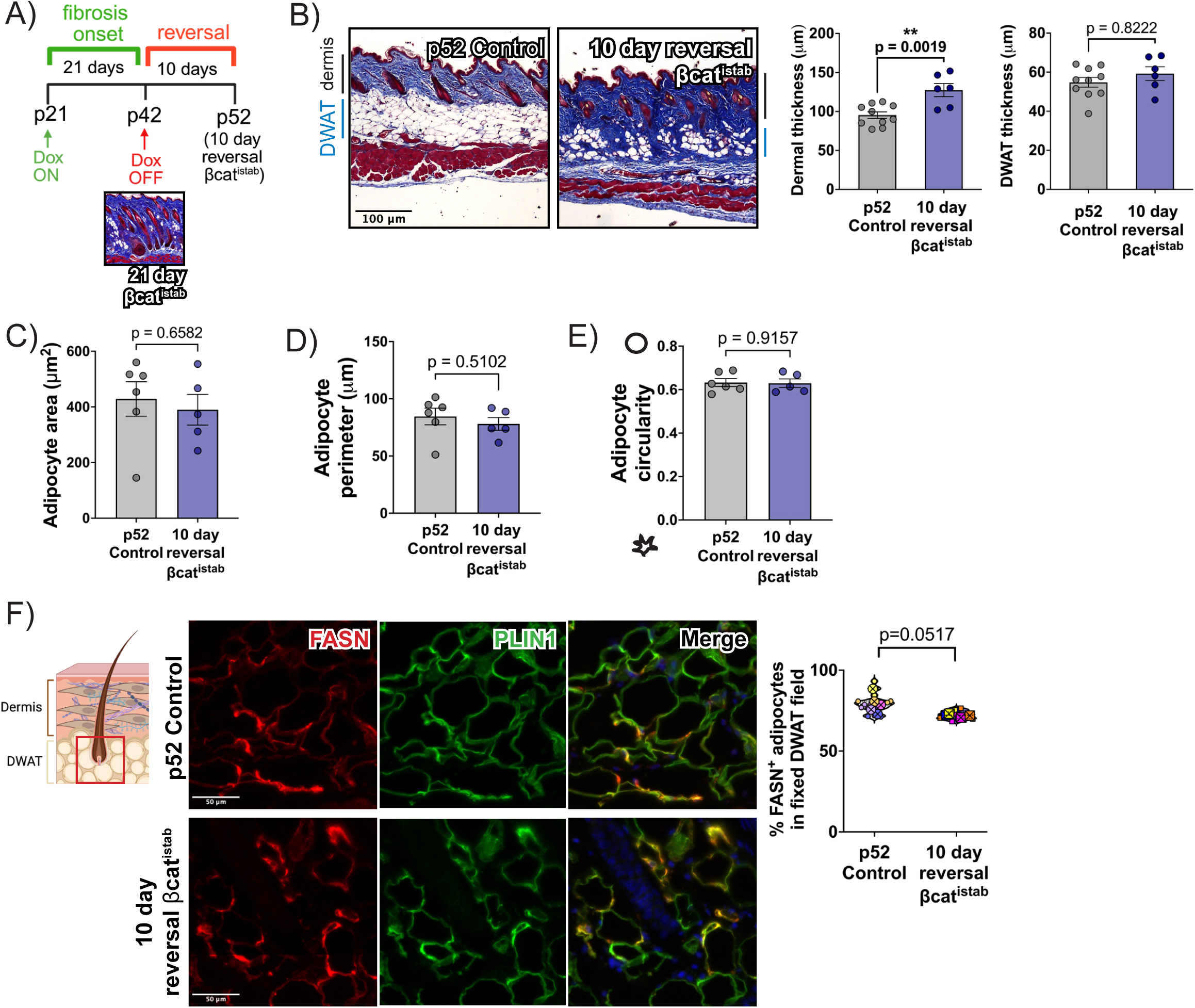
FASN recovers after 10 days of reversal from established fibrosis. A) Schematic representing design of 10-day reversal βcat^istab^ experiment. Inset picture of established fibrosis at 21 days βcat^istab^. B) Histology of dorsal skin of 10-day reversal βcat^istab^ and age matched control and corresponding quantification of dermal and DWAT thickness (n=6-10). C) Quantification of individual adipocyte area, D) perimeter and E) circularity in both groups (n=5-6). F) Immunofluorescence for FASN and Perilipin1 in control and 10-day βcat^istab^ reversal skin and respective quantification of % FASN^+^ adipocytes (n=4-5). P-values were obtained using unpaired students t-test with Welch’s correction for bar and scatter plots. *, **, ***, **** is p-value <0.05, 0.01, 0.001 and 0.0001 respectively. A p-value < 0.05 is considered significant.

### 3.4 Recovery from Wnt-induced lipodystrophy and fibrotic ECM accumulation is dependent on FASN mediated *de-novo* lipogenesis

Since *FASN* expression is suppressed in human fibrotic skin and restored following Wnt pathway inhibition in SSC **(Figure 1H-J)**, we reasoned that FASN-dependent DNL might be functionally required for recovery from skin fibrosis. To test this, we pharmacologically inhibited FASN enzyme catalysis using TVB3664 (FASN_i_) during the reversal phase of the βcat^istab^ mouse model, when Wnt signaling is withdrawn and DWAT and dermal architecture normally recover (**Figure 5A**). FASN_i_-treated mice maintained normal body weight relative to untreated, and vehicle treated controls **(Supplementary** Figure 7A), but exhibited significantly thinner DWAT (**Figure 5B, C)**. Interestingly, the total skin thickness of FASN_i_ mice was significantly lower when compared to vehicle-treated mice **(Supplementary figure S7B)**. Subsequent morphological analysis of adipocytes showed that FASN_i_ mouse skin had smaller and less circular dermal adipocytes **(Figure 5D, E, Supplementary Figure S7C, S7D).**

**Figure 5:**
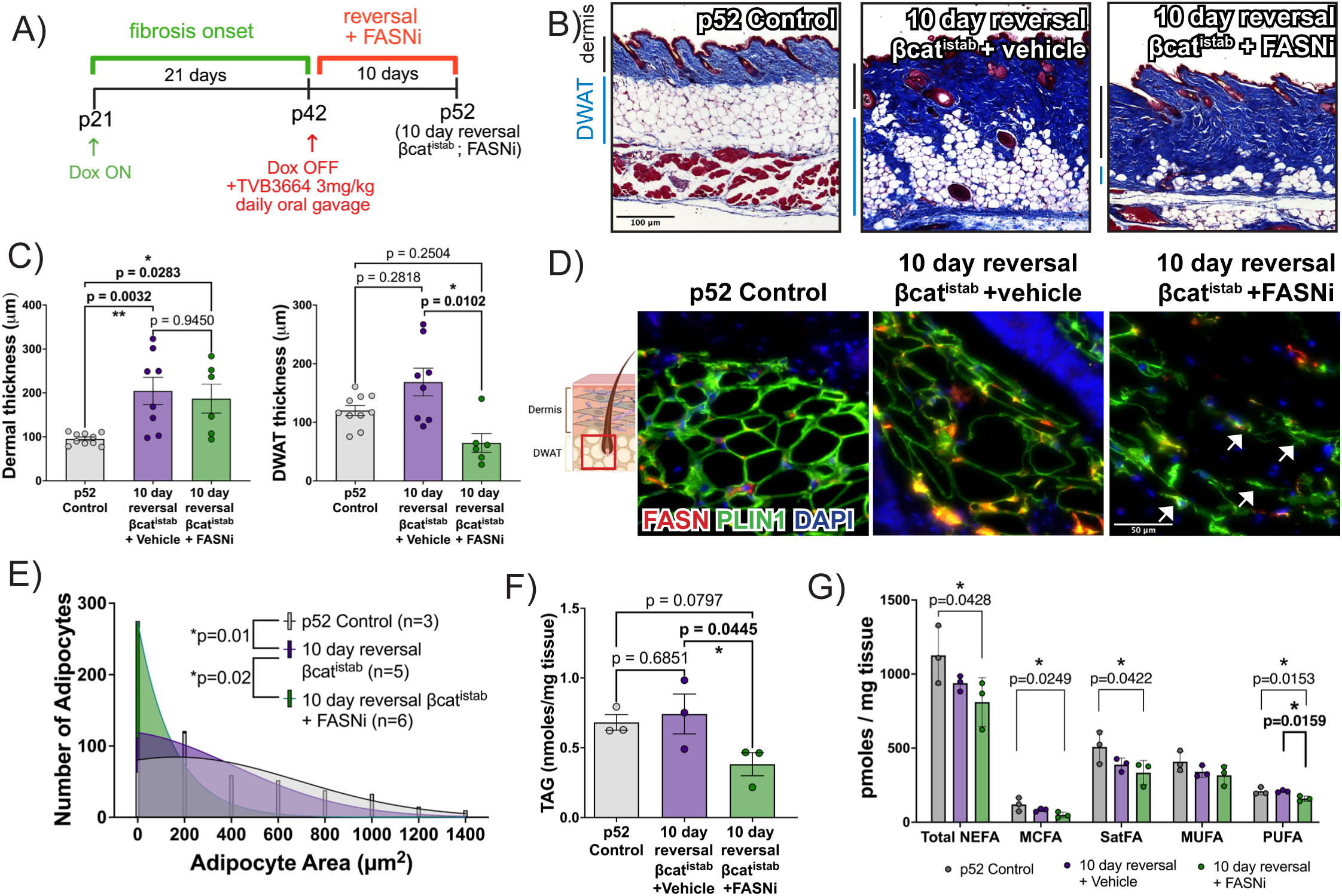
FASN is functionally required for dermal adipocyte refilling after fibrotic lipodystrophy. A) Schematic of regimen for FASN inhibitor treatment. During reversal, mice were given 3mg/mL TVB3664 every day for 10 days by oral gavage. B) Histology of healthy p52 mouse dorsal skin, 10-day reversal βcat^istab^ mouse skin with vehicle gavage and FASN-inhibited 10-day reversal βcat^istab^ mouse skin stained with Massons Trichrome (scale bar = 100μm) C) Dermal and DWAT thickness measurements as absolute values (n=5-7). D) Immunostaining for FASN (red) and Perilipin1 (green) on mouse skin. Arrows point to lipid depleted dermal adipocytes with functionally inhibited FASN protein (scale bar = 50μm). E) Histogram of adipocyte area in all three groups (n=5-6). F) Triglyceride quantity measurements normalized to weight of tissue sample G) Lipidomics analysis of non-esterified fatty acids (NEFA) grouped into by total NEFA, medium chain FA (MCFA), mono-unsaturated FA (MUFA) and poly-unsaturated FA (PUFA) normalized to weight of tissue (n=3) from whole skin of p52 control, 10-day reversal βcat^istab^ mouse skin with vehicle gavage and FASN-inhibited 10-day reversal βcat^istab^ (n=3). P-values were determined by Ordinary 1-way ANOVA with appropriate posthoc test. For the frequency distribution, non-parametric Kruskal-Wallis test was applied. *, **, ***, **** is p-value <0.05, 0.01, 0.001 and 0.0001 respectively. A p-value < 0.05 is considered significant.

Biochemical analysis supported this morphological impairment: FASN_i_-treated skin had significantly lower triglyceride levels, normalized to tissue weight **(Figure 5F)**. Targeted lipidomics of the whole skin NEFA fraction revealed marked decreases of polyunsaturated fatty acids (PUFAs) in the FASN_i_ group compared to vehicle control, while the quantitative sums for Total FA, saturated FA, medium chain FA, and monounsaturated FA (MUFA) remained unchanged (**Figure 5G, Supplementary Figure S7E, S7F, Supplementary table 2).** These data indicate that FASN enzyme function is required, in part, for dermal adipocyte refilling, improvement of adipocyte morphology and NEFA levels during recovery from established fibrosis associated lipodystrophy.

The histology and significantly thicker dermis relative to total skin of FASN_i_-treated mice compared to the vehicle suggested excess collagen matrix in the dermis that accompanied the delay in DWAT recovery **(Figure 6A)**. We found the percent area occupied by collagen was higher in the dermis and DWAT of the FASN_i_-reversal skin compared to the vehicle group **(Figure 6B)**. Topography of collagen and excess sulfated proteoglycan are also indicators of skin fibrosis and recovery (17,51,52). We examined the topography of the collagen matrix and abundance of sulfated proteoglycans in the dermis and DWAT in FASN_i_-reversal skin. Mouse dorsal skin sections were stained with Picosiris red (PSR) and imaged with polarized light to reveal birefringent collagen fibers in the skin. The Workflow Of Matrix BioLogy Informatics (TWOMBLI) was utilized to quantify different collagen fiber properties in control, vehicle gavaged, and FASN-inhibited reversal skin (52,53) **(Figure 6C)**.

**Figure 6:**
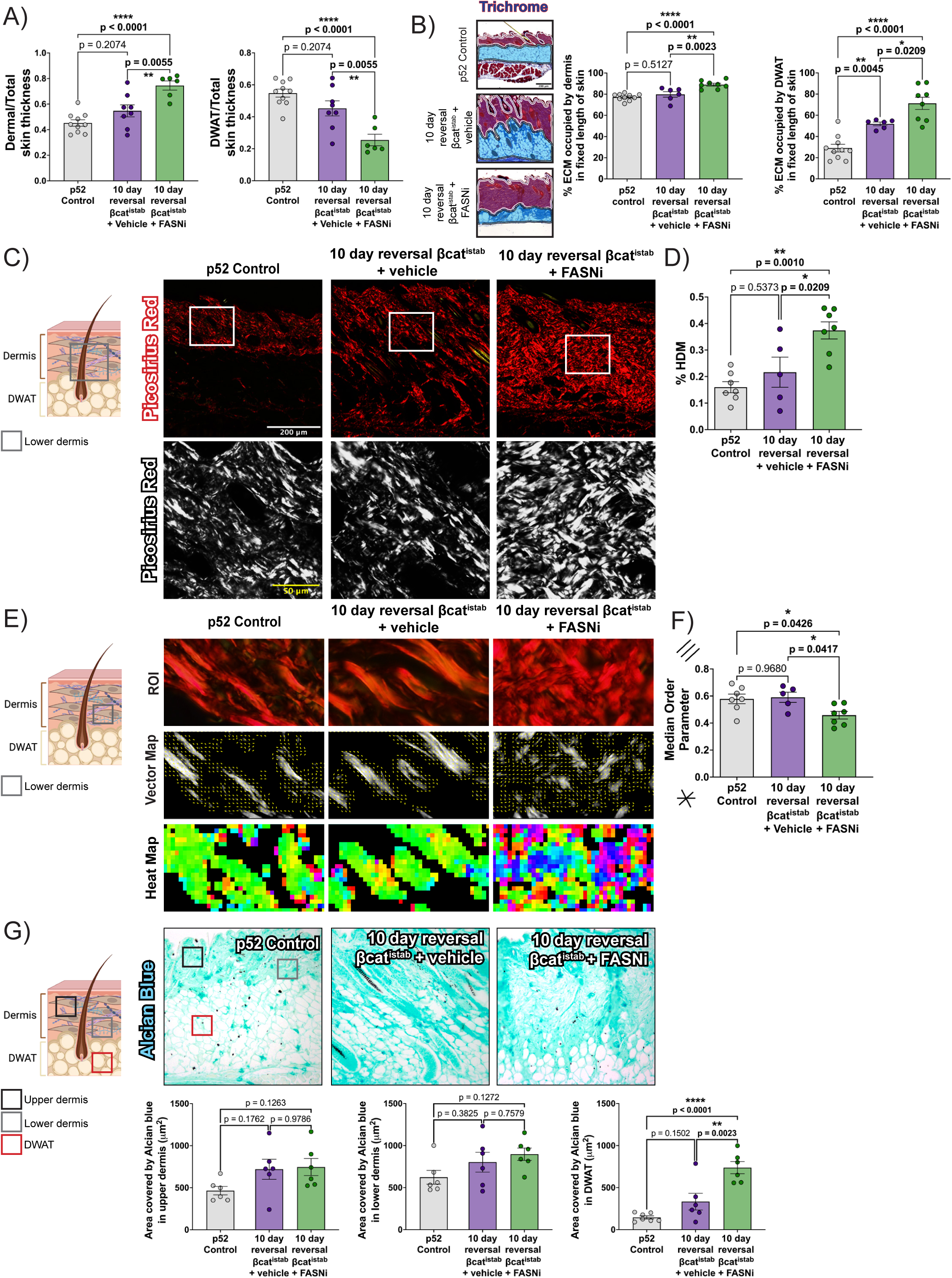
FASN is required for resolving fibrotic matrix during recovery from Wnt-induced fibrosis. A) Dermal and DWAT thickness as a ratio over total skin thickness in p52 mouse dorsal skin, 10-day reversal βcat^istab^ mouse skin with vehicle gavage and FASN-inhibited 10-day reversal βcat^istab^ mouse skin (n=6-10). B) Massons trichrome stained mouse skin outlining dermis (red with white border) and DWAT (blue with black border) of all three groups and respective quantification of percentage of ECM occupied by dermis and DWAT (n=6-10). C) Collagen was stained using Picosiris red (PSR) and visualized with polarized light microscopy in all three groups. Insets show high magnification of collagen fibers in black and white. D) Quantification of percent high density matrix in all three groups using image analysis algorithm TWOMBLI (n=5-7). E) Cropped ROIs from PSR-stained polarized light 40X images (top row), and corresponding vector (second row) and heat maps (third row) showing orientation of collagen using Alignment by Fourier Transform (AFT) algorithm. F) Quantification of median order parameters calculated from the vector maps for all three groups (n=5-7). G) Alcian blue (at pH2.5) stained skin to detect sulfated proteoglycans and respective quantification for area occupied by sulfated proteoglycans in upper dermis, lower dermis and DWAT of skin from all three groups. P-values were obtained by one-way ANOVA with Tukey’s multiple comparisons for equal variances or Brown Forsythe and Welch ANOVA with Dunnett’s T3 multiple comparisons when variances are unequal. *, **, ***, **** is p-value <0.05, 0.01, 0.001 and 0.0001 respectively. A p-value < 0.05 is considered significant.

Principal Component analysis of all fiber properties from the lower dermis of all three groups showed that the FASN_i_-reversal skin is distinct from the other two groups, driven by High density matrix (HDM) **(Supplementary Figure S8A)**. HDM, which describes the amount of matrix in a fixed space, and alignment of matrix are two crucial fiber properties involved in fibrosis onset and recovery (52,53) **(Figure 6C)**. Percent of HDM, but not alignment, was significantly higher in FASN_i_ reversal skin compared to vehicle-treated controls (**Figure 6D, Supplementary Figure S8B)**. Alignment by Fourier Transform (AFT), an independent method to quantify the alignment of collagen fibers, shows that collagen alignment is significantly decreased in FASN_i_-reversal skin (52,54) **(Figure 6E, F)**. The area occupied by sulfated dermal proteoglycans (stained with Alican blue at pH 2.5), but not highly acidic sulfated proteoglycans (stained with Alican blue at pH 1.0), are significantly increased in FASN_i_ reversal skin **(Figure 6G, Supplementary Figure S8C)**. All together these data show that FASN_i_ hampers lipogenesis in adipocytes during reversal and subsequently impedes the recovery from fibrotic ECM.

Collectively, these results demonstrate that FASN activity is essential for restoring adipocyte morphology, lipid content, and matrix resolution during recovery from Wnt-induced fibrosis. We propose that properly functioning adipocytes contribute to ECM remodeling via DNL-mediated metabolic and structural outputs. This mechanism likely underlies the therapeutic effects of Wnt inhibition (e.g., C-82) in human SSc skin, where restoration of adipocyte function may drive fibrosis resolution. Our working model highlights the Wnt–DNL–FASN axis in adipocytes as a central regulator of both lipodystrophy onset and fibrosis recovery **(Figure 7).**

**Figure 7:**
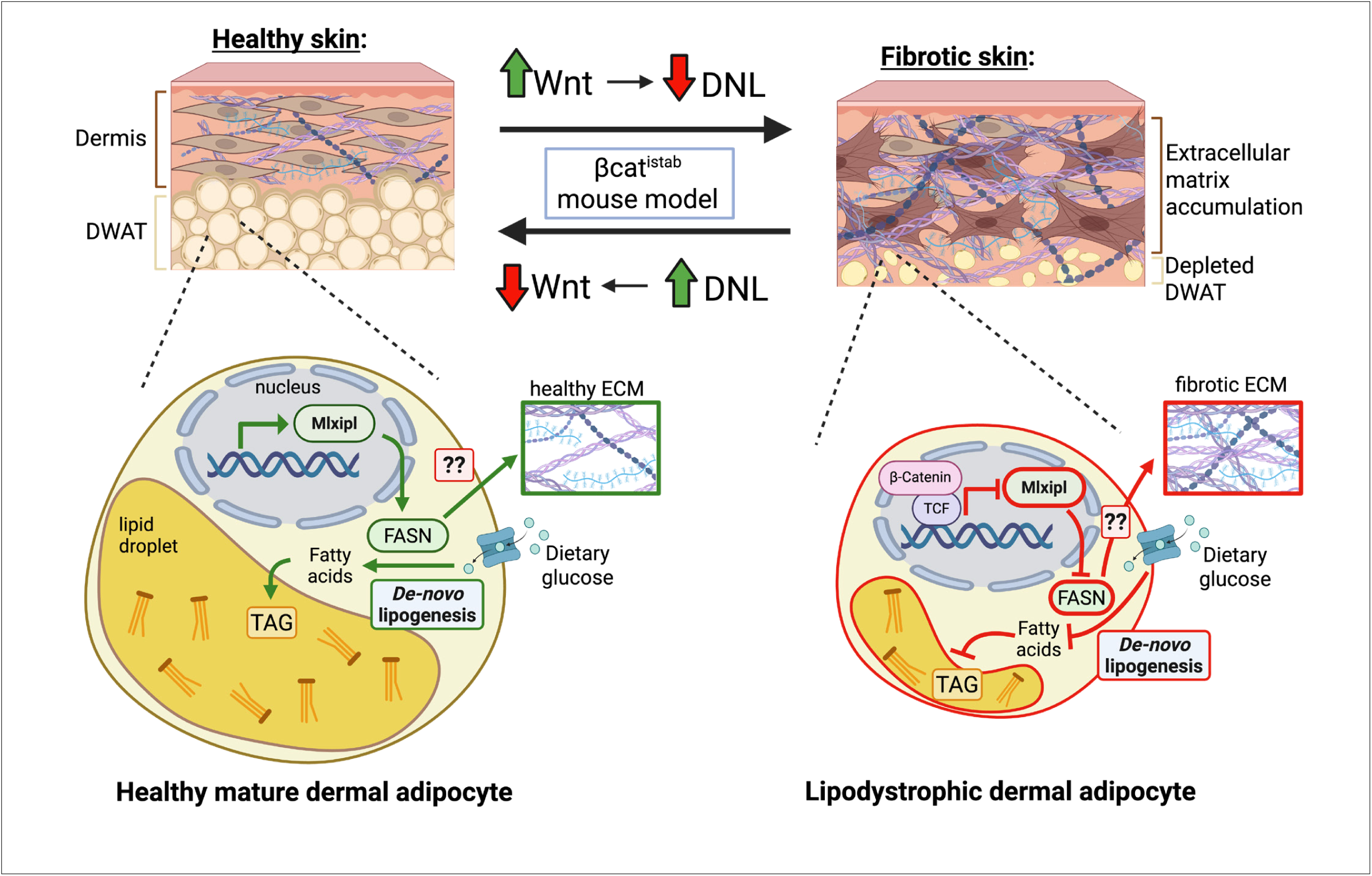
Proposed working model of Wnt-induced downregulation of *de-novo* lipogenesis in mature dermal adipocytes during skin fibrosis. In healthy skin, dermal adipocytes undergo *Mlxipl*/FASN-mediated *de-novo* lipogenesis to generate *de-novo* fatty acids which enters the lipid droplet, leading to big lipid-filled mature adipocytes and homeostatic ECM production in the dermis. In fibrotic skin, over-expression of β-catenin transcriptionally downregulates *Mlxipl*/FASN-mediated *de-novo* lipogenesis, leading to small lipid-depleted fibrotic adipocytes and fibrotic ECM production in the dermis. *Mlxipl* – Mlx interacting protein like (encodes for ChREBP), FASN – Fatty acid synthase, ECM – extracellular matrix, DNL – *de-novo* lipogenesis, TAG – triacylglycerides, DWAT – dermal white adipose tissue. Created in https://BioRender.com.

## 4. DISCUSSION

Acquired lipodystrophy can arise from diverse causes and affects tissue function across multiple organs (25), including complex etiologies of tissue fibrosis and the associated lipodystrophy in skin, lung, and heart (5,55). While current research and clinical strategies largely focus on immune and extracellular matrix dysregulation, the therapeutic efficacy remains limited (56–58). This underscores the need to identify alternative mechanisms that control adipocyte function and tissue remodeling during acquired lipodystrophy and fibrosis. Here we identify suppression of *de-novo* lipogenesis (DNL) as an early and reversible event in dermal lipodystrophy and skin fibrosis, downstream of sustained Wnt signaling in mature dermal adipocytes. Using a genetically inducible and reversible model of Wnt pathway activation in dermal adipocytes, we demonstrate that transcriptional downregulation of FASN and other DNL-axis enzymes is Wnt-dependent and cell autonomous *in vivo*. Moreover, FASN downregulation emerges as an early feature in human fibrosing conditions such as SSc and keloids, as well as in Wnt-induced models of fibrotic lipodystrophy. Importantly, we show that FASN-dependent FA synthesis is required for refilling dermal adipocyte triglycerides and resolving fibrotic ECM during recovery from Wnt-induced fibrosis **(Figures 5,6)**. Together, these findings establish a mechanistic link between Wnt activation of adipocyte FA catalysis and ECM remodeling and suggest that therapeutic strategies aimed at restoring local DNL in dermal adipocytes may provide a new approach to mitigate lipodystrophy-associated fibrosis.

Fatty acids have been implicated in fibrosis either as fibro-protective or profibrotic mediators, depending on the tissue-specific context and the fibrotic manipulation engaged *in vivo*. For example, *in vitro* treatment of fibroblasts with 10uM palmitic acid decreased cell proliferation and downregulated collagen 1 (*Col1*) and Elastin (*Eln*) mRNA expression (59), indicating an anti-fibrotic effect on ECM components. In a different context, dermal fibroblasts in a radiation-induced skin fibrosis model *in vivo* exhibit an anabolic metabolic phenotype marked by increased glycolysis and suppressed FA oxidation (FAO) to support ECM protein production like Collagen and Vimentin (20). In this model, upregulation of FAO via FA transporter CD36 by dermal fibroblasts promoted resolution of fibrosis, however, the source and identity of FA taken up by CD36^hi^ fibroblasts remains undefined (20). In bleomycin and Wnt-induced models of skin fibrosis, DWAT lipodystrophy occurs via ATGL-dependent lipolysis, and the resulting secretome of lipids and non-esterified “Free” FA (FFA) products has been proposed to influence fibrotic ECM expansion in different ways (5,16–18,60). On the one hand, systemic pharmacological inhibition of ATGL using atglistatin or genetic ATGL knockout in bleomycin treated mice preserves DWAT but also enhances dermal thickness and collagen deposition, suggesting that FFA may act to constrain ECM accumulation (16,60). However, in our tissue-specific Wnt activation model, we found that ATGL-dependent lipolysis in the dermal adipocytes was required for fibrotic ECM expansion and remodeling in the dermis, indicating that dermal adipocyte-derived FFAs may also promote ECM accumulation depending on the context (18). Together, these seemingly contradictory findings point to a complex and context-dependent role of FA, lipid metabolism, and fibrotic ECM expansion, highlighting the need to further dissect distinct lipid metabolic pathways and identify lipid specific profiles that can mitigate poor fibrotic outcomes.

The DNL pathway is transcriptionally regulated by two well-characterized metabolic transcription factors, the carbohydrate responsive *MLXIPL/*ChREBP or *SREBF1*/SREBP1 activated downstream of insulin signaling. Whether Wnt signaling regulates these master transcription factors may depend on the experimental model and tissue context of Wnt activation. For instance, Wnt pathway inhibition in adipocytes, isolated from the outer ear and differentiated *in*-*vitro*, led to downregulation of DNL axis genes and transcriptional regulators, and this repression could be compensated by other cell types present in the SVF fraction (39). In contrast, adipocyte-specific deletion of CTNNb1 transcriptional co-factor TCF7L2 results in activation of the DNL program and adipocyte hypertrophy in inguinal WAT (40). Consistent with this, Wnt pathway activation in *Drosophila* larval adipocytes leads to downregulation of DNL genes and regulators (61). Our current results align with these gain-of-function models, supporting that Wnt activation is sufficient to suppress the DNL transcriptional program in dermal adipocytes, however, whether Wnt activation is required to downregulate DNL axis and its regulators in dermal adipocytes remains unresolved. Our meta-analysis suggests that *MLXIPL* (ChREBP) may be a direct transcriptional target of Wnt/β-catenin action, supported by the TCF7L2 ChIP-seq peaks at the *MLXIPL* locus in other contexts. Future work will be needed to define how Wnt signaling can remodel chromatin accessibility and recruit transcriptional co-repressors to suppress the DNL program in dermal adipocytes.

In fibrotic skin diseases such as SSc, morphea, keloids and chronic wounds, the dermal adipose layer is frequently atrophied and replaced by ECM-rich fibroblasts (62). Gene profiling studies of human SSC fibrotic skin have consistently identified lipid metabolic pathways as dysregulated, particularly for FA metabolism, lipid biosynthesis, and lipolysis (16,18,32,33,63). However, most of these studies did not stratify samples by disease duration, making it unclear whether changes in lipid metabolism are drivers or consequences of fibrosis. In contrast, our findings support a causative role for early suppression of DNL in dermal adipocytes during fibrotic onset. Specifically, we show that Wnt activation leads to early downregulation of FASN and DNL axis genes, consistent with the decreased *FASN* mRNA expression in SSc and keloid skin. Supporting this, Burja et al., recently identified fatty acid synthesis via FASN as one of the most consistently downregulated lipid metabolism processes in SSc skin (63). Our results align well with studies demonstrating reactivation of FASN and DNL axis gene expression following topical administration of Wnt inhibitor C-82 in SSc patients (46), and with improved FASN expression in SSc patients treated with mycophenolate mofetil (MMF) (32).

While our model provides strong evidence that Wnt activation in mature dermal adipocytes suppresses DNL and impairs adipocyte recovery, the causal intermediates (e.g., specific lipid species or co-repressors of ChREBP/SREBP1) remain undefined. We focused on mature dermal adipocytes, however, prior work from our lab and others has shown that Wnt-CTNNB1 signaling regulates fibrotic gene expression in adipocyte precursor and stem-like cells (64,65). Future studies will need to investigate the extent to which early suppression of DNL in adipocyte precursors vs. mature adipocytes contributes to fibrosis across disease stages, metabolic phenotypes like obesity, or tissue niches. Additionally, due to the absence of DNL or FASN agonism tools, we were limited in our capacity to activate DNL during fibrosis in the Wnt activated mouse model. Lastly, our targeted lipidomic analysis was performed at the whole-skin level; future work using unbiased or cell-type resolved lipidomics will be essential to attribute multiple lipid changes to specific tissue compartments or cellular states.

Despite the lack of clarity on precise mechanisms, emerging regenerative therapies such as fat grafting, adipocyte stem cell treatment, extracellular vesicles (EVs), and exosomes, have shown promise for treating fibrotic skin conditions (66–73). Our study adds mechanistic insight showing that pharmacological inhibition of FASN leads to a decrease in omega-6 and omega-3 PUFA in whole skin, which coincides with increased ECM deposition in both dermis and DWAT. This finding is consistent with prior studies showing that FASN inhibition worsens bleomycin-induced lung fibrosis in mice, while fatty acid supplementation or LXR-α agonists show protective potential as treatments of lung fibrosis (74–76). In conclusion, future studies are needed to define the specific lipid species and lipid metabolic regulators that can restore adipocyte function and halt fibrotic progression, thereby harnessing the therapeutic potential of FASN and the DNL pathway for treating fibrotic skin disease.

## Author Contributions

All authors were involved in drafting the article or revising it critically for important intellectual content, and all authors approved the final version to be published. RPA had full access to all the data in the study and takes responsibility for the integrity of the data and the accuracy of the data analysis. Study Conception and Design: RPA, MCR and SRM. Acquisition of Data: SRM, PM, RV, MM, QM, KK, RP. Analysis and Interpretation of Data: RPA, MCR, SRM, PM, RV, MM. Writing and editing: RPA, MCR and SRM wrote the manuscript which was edited by all the authors. All the authors have approved the submitted version.

## Acknowledgments

We thank all the past and present members of the Atit lab for their input and feedback on this project. We thank Morgan Horowitz for quantification of Picrosirius red staining using TWOMBLI, Rachel Kim and Emilia Sanz-Rios for quantification of Trichrome staining, Megan Gregory for genotyping the mice, and Saada Eid for technical support with the bleomycin mouse model. Special thanks to Anna Jussila, Valarie Horsley, David Buchner, Timothy Mead, Rodrigo Somoza-Palacios and William Spencer for technical support and advice. We thank the bio shared instrumental facility, Tissue resources core and Genomics core facility at Case Western Reserve University and CWRU-ENGAGE program for summer fellowships (Moore). This project was funded by National Institutes of Health, National Institute of Arthritis and Musculoskeletal and Skin Diseases R01 AR076938 (Radhika Atit and Valerie Horsley) and National Scleroderma Foundation (Suneeti Madhavan and Qiannan Ma).

## Data Availability

Sequencing and analysis data were deposited in the National Center for Biotechnology Information Gene Expression Omnibus under accession number GSE305616 (mouse dermal adipocyte bulk RNA sequencing), GSE130995 (human SSc RNA sequencing data), GSE90051 (keloid microarray data) and GSE94340 (human C-82 clinical trial microarray data. Data will be made available after publication and upon request.

