## Supplementary material for "Lipodystrophy and recovery are mediated by the Wnt/lipogenesis axis during skin fibrosis"

**Supplementary Methods:**

*Intradermal adipocyte isolation from whole skin*

Fresh dorsal dermis was collected, minced and incubated in a digestion solution containing 2 mg/mL Type I collagenase (Worthington Biochemicals, LS004196), 2.5mM glucose (Sigma-Aldrich, G8769), 3% BSA (Fisher Scientific, BP1600) and 200nM adenosine (Sigma Aldrich, A9251) in Krebs-Ringer-HEPES (KRH) buffer. After 45 minutes of incubation with constant rotation, the digest was spun down at low speed (400xg, 15 minutes) and the top floating adipocyte fraction was collected. The cells were washed briefly in KRH buffer and stored in TRIzol reagent (Thermo Fisher Scientific, 15596026). The stromal fraction was used as a control for RNASeq validation studies.

*DWAT biopsy collection*

A large piece (2cmx2cm) of dorsal skin from mice were collected after dissection after cut into smaller 2mm biopsy pieces. Using blunt forceps to hold the edge of the tissue, a scalpel was used to scrape the bottom of the tissue

*In vivo whole skin lipidomics and triglyceride assay*

Mouse skin was harvested and flash frozen immediately during dissection and whole skin lipidomics was performed using a previously described protocol [77]. Briefly, skin punches, 60-90mg in weight, were homogenized in 1ml 0.1M potassium phosphate buffer, pH 6.8, using a bead mill (Fisher Scientific, Bead Mill 24) and 1.4mm ceramic beads (Fisher Scientific, 15-340-159). The lysates were transferred to 13x100mm borosilicate glass culture tubes (Fisher Scientific, #14-961-27). For triglyceride assay, 2ul of the lysate was used for spectrophotometry analysis using TG Infinity TAG reagent (Fisher, TR22421) at 540 nM. Amount of triglyceride (in moles) was obtained using a glycerol standard curve and normalized to tissue weight. Lysates were then acidified with 40ul of prewashed (with Hexanes) 1N HCl, vortexed briefly, and 1ml methanol was added to the tubes and vortexed again. Total lipids were extracted twice by adding 2ml of Hexanes; for each extraction, samples were vortexed vigorously for 10-15 sec and centrifuged at 2000g for 1 minute to complete phase separation. The upper organic layer from each extraction was combined into a new 13x100 mm borosilicate glass tube. Extracted lipids were brought to dryness and resuspend in 300ul 2,2,4-Trimethylpentane (isooctane). For the unlabeled non-esterified fatty acid (NEFA) fraction, 50ul of the resuspended total lipids was transferred to a new borosilicate glass tube, mixed with (25ng) blended stable isotope internal standard, taken to dryness under gaseous N2, and resuspended in 25ul of 1% pentafluorobenzyl bromide in acetonitrile to which 25ul of 1% diisopropylethylamine in acetonitrile was added and samples were incubated at room temperature for 30 minutes. Pentafluorbenzyl-fatty acid derivatives were taken to dryness under gaseous N2 and resuspended in 100ul hexanes. For the unlabeled total fatty acid fraction (TFA), 20ul of the original 300ul total lipid extract was transferred to a separate Teflon lined screw cap glass tube, mixed with (75ng) blended stable isotope internal standard, and taken to dryness under gaseous N2. TFA samples were resuspended in 500ul of 100% ethanol to which 500ul of 1M NaOH was added to saponify the TFA fraction at 90C for 30 minutes, followed by acidification using 550ul of 1M HCl. Saponified samples were then extracted twice with 1.5ml of Hexanes, taken to dryness under gaseous N2, and derivatized as above. Derivatized TFA samples were resuspended in 300ul hexanes for injection. For the NEFA and TFA fractions, 1ul of pentafluorbenzyl-fatty acid derivatives was injected and data were collected on the GC-MS (8890 GC, 5977B MSD, Agilent) DB-1MS UI column (122-0112UI, Agilent) with the following run program: 80C hold for 3 minutes, 30C/minute ramp to 125C, no hold, 25C ramp to 320C and hold for 2 minutes. The flow for the methane carrier gas was set at 1.5ml/minute. Data were acquired in full scan negative chemical ionization mode to identify fatty acids of acyl chain length from 8 to 22 carbons. Peak areas of the analyte or of the standard were measured, and the ratio of the area from the analyte-derived ion to that from the internal standard was calculated.

*Quantitative PCR*

DWAT biopsy samples or cell culture samples were collected in TRIZol. For DWAT biopsies, tissue was homogenized using a motor and pestle. RNA was extracted using the manufacturer's protocol. RNA concentration was determined using ThermoFisher NanoDrop (8000TS). Reverse transcriptase-PCR was performed using MultiScribe Reverse Transcriptase (Thermo, 4311235). Qualitative PCR (qPCR) was performed in duplicate on 4ng of cDNA per reaction with dye-based (FAM) Taqman probes (ThermoFisher Scientific, 4331182) for *Plin1*, *Axin2*, *Acly*, *Acaca, Fasn*, *Slc2a4* (Glut4)*, Mlxipl* and *Srebf1* along with *Hprt* or *ActB*, as the housekeeping control. Comparative $\Delta\Delta C_T$ method was used for obtaining relative qualification of each gene with respect to the housekeeping and internal control (no Wnt activation).

*Image analysis for adipocyte counts and size*

Adipocytes were manually counted using the FIJI cell counter in 40X merged images of dorsal skin sections co-stained with FASN-Plin1 or FASN-ConA. Adipocytes counted were segregated into cells which contained FASN expression and cells that did not have FASN. For adipocyte area measurements, cells were manually circled using the FIJI segmentation editor tool and analyzed for area (in micron squared).

*FASN corrected fluorescence analysis*

Corrected fluorescence intensity was calculated from immunofluorescence images using Fiji/ImageJ. The DWAT of PBS-treated and BLM-treated mice were manually traced to obtain DWAT ROIs from the single channel FASN images. For fluorescence intensity, images were transformed to 8-bit and the average of mean gray values of three circles on non-fluorescence regions was calculated as the background fluorescence using standard methods [75]. The corrected fluorescence per ROI was calculated as integrated density – (area of ROI × average of background fluorescence).

*Masson's Trichrome staining and quantification*

Formalin fixed paraffin embedded (FFPE) mouse skin were stained with Masson trichrome using CWRU Tissue resources core. Detection of collagen area occupied by blue was performed using FIJI/ImageG by color thresholding Hue (135-195), Saturation (0-25) and Brightness (0-20) and converting to binary. ROIs of the dermis and DWAT were manually traced and the percentage of area occupied by collagen in white was measured.

*Picrosirius red staining and quantification*

FFPE mouse skin was stained with picrosirius red (26357–02; Electron Microscopy Science) for 20 minutes and imaged with polarized light to detect bifringent collagen fibers as described before [18,50]. Image analysis using TWOMBLI and AFT algorithms were performed as previously described [18,50]

*PCA (Principal Component Analysis)*

PCA was comprised of TWOMBLI output and the covariance between variables was checked in RStüdio. Each data point is one animal with outliers removed before analysis as calculated by GraphPad Prism. PCA biplot was generated by principal components 1 and principal component 2.

*Alcian Blue staining and quantification*

Alcian Blue staining at pH 2.5 and pH 1.0 was undertaken using protocols adapted from Lai, M. & Lü, B [76]. Alcian Blue at pH 2.5 and 1.0 allowed visualization of the expression of widely distributed and highly acidic sulfated proteoglycans, respectively. ROIs from the upper dermis, lower dermis, and DWAT were used and run through a FIJI/ImageJ macro with color threshold parameters (Hue: 0-255, Saturation: 75-255, Brightness: 0-200) before measuring the area covered by Alcian blue.

*Adipocyte cell culture*

Stromal vascular fraction (SVF) from collagenase skin digest was filtered and spun down to collect isolated SVF cells which were then plated, cultured and passaged 2-3 times till 80% confluence. Cells were differentiated into adipocytes using adipocyte induction media (AIM) (DMEM (Thermofisher, 11995065) containing glucose, pyruvate, 10% fetal bovine serum (FBS, Sigma Aldrich, F0926), 100µM indomethacin, 1 µM dexamethasone, 500 µM 3 -isobutyl-1-methylxanathine (IBMX), and 10µM insulin) for 8-12 days, with a media change every 2$^{nd}$ day. Lipid filled adipocytes were then kept in adipocyte maintenance media (AMM) (DMEM with 10% FBS and 10µM insulin). For Wnt activation, replicate cultures were treated with either 7µM CHIR99021 (Cayman, 13122) or Wnt agonist BML284 (Cayman, 19903), with media prepared fresh and changed every day.

*In vitro lipid staining*

After 5 and 8 days of CHIR treatment, cells were rinsed and fixed in 10% buffered formalin for staining. Cells were stained with Oil red O (ORO) (Sigma Aldrich, O0625-25G) for LipidSpot™ A488 (Biotium 70065) for 10 minutes and rinsed with PBS. For LipidSpot™ staining, cultures were co-stained with DAPI (4',6-diamidino-2-phenylindole) before being imaged on a Leica DMi8 inverted microscope on green and blue filters. Each well was photographed in 2 non-overlapping regions.

*In vitro lipid area and droplet analysis*

Oil red staining was analyzed by Cell Profiler software on 40X fixed field as previously described (A. R. Jussila et al., 2022). Area covered by LipidSpot™ was measured using FIJI/ImageJ and normalized to the number of nuclei in the fixed field. Briefly, green (for LipidSpot™) and blue (DAPI) channels underwent a gaussian blur (sigma=2,3 respectively) followed by thresholding and converting the image into a binary mask. Area occupied was measured for the green channel while the number of particles (size criteria 30-infinity pixels) was analyzed for the blue channel. For individual droplet analysis, LipidSpot™ stained droplets were segmented using CellPose and fed into FIJI/ImageJ to extract droplet measurements such as area, perimeter and circularity. Images were automatically segmented by a cellpose custom trained model. The custom trained human-in-loop model was created by training cyto3 pretrained cellpose model, on the ROIs of 5 images segmented by a combination of cyto3 and hand editing [79].

*In vitro $^{13}$C acetate labeling, lipidomics, tracer studies and triglyceride assays*

Lipidomics and tracing studies were performed in Dr. Michael Rudolph's lab at the University of Oklahoma Health Sciences Center (OHSU) as previously described [40,77]. Briefly, cells were lysed in 1ml 50% methanol (in HBSS buffer). The lysates were transferred to 13x100mm borosilicate glass culture tubes (Fisher Scientific, #14-961-27). Lysates were acidified with 10ul of prewashed (with Hexanes) 1N HCl and vortexed briefly. Total lipids were extracted twice by adding 1ml of isooctane:ethyl acetate 3:1 (vol/vol) and once with 1ml Hexanes; for each extraction, samples were vortexed vigorously for 10-15 sec and centrifuged at 2000g for 1 minute to complete phase separation. The upper organic layer from each extraction was combined into a new 13x100 mm borosilicate glass tube. Extracted lipids were brought to dryness and resuspend in 300ul 2,2,4-Trimethylpentane (isooctane). For triglyceride assay, 30ul of resuspended total lipids was used for spectrophotometry analysis using TG Infinity TAG reagent (Fisher, TR22421) at 540 nM. Amount of triglyceride (in moles) was obtained using a glycerol standard curve and data was analyzed For the unlabeled non-esterified fatty acid (NEFA) fraction, 100ul of the resuspended total lipids was transferred to a new borosilicate glass tube, mixed with (25 ng) blended stable isotope internal standard, taken to dryness under gaseous N2, and resuspended in 25ul of 1% pentafluorobenzyl bromide in acetonitrile to which 25ul of 1% diisopropylethylamine in acetonitrile was added and samples were incubated at room temperature for 30 minutes. Pentafluorbenzyl-fatty acid derivatives were taken to dryness under gaseous N2 and resuspended in 100ul hexanes. For the unlabeled total fatty acid fraction (TFA), 50ul of the original 300 ul total lipid extract was transferred to a separate Teflon lined screw cap glass tube, mixed with (66.7 ng) blended stable isotope internal standard, and taken to dryness under gaseous N2. TFA samples were resuspended in 500ul of 100% ethanol to which 500ul of 1M NaOH was added to saponify the TFA fraction at 90C for 30 minutes, followed by acidification using 550ul of 1M HCl. Saponified samples were then extracted twice with 1.5ml of Hexanes, taken to dryness under gaseous N2, and derivatized as above. Derivatized TFA samples were resuspended in 267ul hexanes for injection. For 13C-2 acetate tracer incorporation analysis, 50ul of the resuspended total lipids was transferred to a new Teflon lined screw cap glass tube, mixed with (66.7 ng) d31 palmitate internal standard and taken to dryness under gaseous N2. The samples were saponified, extracted and derivatized as above. Derivatized samples were resuspended in 267ul hexanes for injection. For the NEFA, TFA and tracer fractions, 1ul of pentafluorbenzyl-fatty acid derivatives was injected and data were collected on the GC-MS (8890 GC, 5977B MSD, Agilent) DB-1MS UI column (122-0112UI, Agilent) with the following run program: 80C hold for 3 minutes, 30C/minute ramp to 125C, no hold, 25C ramp to 320C and hold for 2 minutes. The flow for the methane carrier gas was set at 1.5ml/minute. Data were acquired in full scan negative chemical ionization mode to identify fatty acids of acyl chain length from 8 to 22 carbons. Peak areas of the analyte or of the standard were measured, and the ratio of the area from the analyte-derived ion to that from the internal standard was calculated. Tracer (13C-2 acetate) incorporation in palmitic acid was analyzed in two carbon increments from 13C-2 to 13C-16 against the d31-palmitate internal standard.

*Western Blot*

Cells were scraped and collected in 1X RIPA (diluted from 10X) (Cell signaling, 9806S). Protein was extracted from samples using a double centrifugation step (to remove maximum lipid content from top layer) and the resulting supernatant was analyzed for protein concentration using the Pierce BCA Protein Assay Kit (Thermo Scientific, PI23227). Protein samples were diluted to 0.5ug/uL in Laemmli buffer (Bio-rad, 1610737) and boiled at 100C for 10 minutes. 10ug of protein was loaded onto a 4-15% SDS-PAGE pre-cast stain-free gel (Bio-rad, 4568084) and run for 60 mins at 250V. The gel was transferred to a PVDF membrane (Bio-rad, 1620174) for 60 mins at 100V. The membrane was imaged using the stain-free blot option in the Bio-Rad ChemiDoc for total protein bands. The membrane was then incubated in blocking buffer (5% milk in TBST) at RT for 1 hour before incubating with rabbit-FASN primary antibody (1:1000, AbCam, ab22759) overnight. Following this, the blot was incubated with 1:10000 goat anti-rabbit peroxidase secondary antibody for 1 hour. Finally, the blot was developed using ECL Western Blotting Detection Reagent (Cytvia, RPN2109) using the manufacturer's protocol. Band intensity and fold change was calculated with respect to total protein content in each lane using ImageLab software.

*Human RNA Sequencing analysis*

Skin biopsies from 33 healthy controls and 48 early-stage (mean disease duration of 1.3 years) SSc patients were analyzed. Patient selection and sample treatments were as described in the PRESS cohort [32]. Briefly, 3- or 4-mm forearm biopsies for each patient were collected and immersed in RNAlater solution, flash frozen and stored in dry ice. miRNeasy Mini kits (Qiagen) were used for RNA extraction and Agilent 2100 Bioanalyzer (Agilent Technologies) was used for RNA integrity testing. Illumina TruSeq stranded Total RNA Library Prep Gold kit was used for cDNA library preparation according to the manufacturer's protocol. Agilent 2200 TapeStation (Agilent Technologies) was used for cDNA quality testing and KAPA Library Quantification Kit (KAPA Biosystems) was used for cDNA quantification before sequencing. 10 pM concentration of the libraries was loaded on cBot (Illumina) and used HiSeq 2500 (Illumina) for a 2 × 76 bp paired-end sequencing. For each sample, 50 million reads were generated. These data are publicly available in the NCBI GEO database under the accession numbers GSE130995.

*Human microarray analysis*

Publicly available microarray datasets GSE90051 (keloid) and GSE94340 (SSc Wnt inhibitor trial) in the NCBI GEO database were mined for differential expression of genes of interest. In GSE90051, biopsies of active keloid lesions and matching adjacent normal skin from seven patients were transcriptionally profiled. Tissue procurement and processing was as described [38]. Briefly, 5mm skin biopsies was flash frozen and homogenized after which tissue RNA was extracted using TRIzol reagent (Invitrogen). Total RNA (1 µg) was amplified by a low RNA input fluorescent linear amp kit (Agilent Technologies) and labeled with cyanin (Cy)3 and Cy5 for control and keloid samples, respectively (CyDye; PerkinElmer, Boston, MA), during the in vitro transcription process, according to the manufacturer's protocol. Hybridization was performed according to the Agilent 60-mer oligo microarray processing protocol using the Gene Expression Hybridization kit (Agilent Technologies). In GSE94340, 26 patients with diffuse SSc (median duration = 8 months) were equally segregated into placebo-treated and Wnt inhibitor treated groups [39]. The study drug and regimen were 0.5% topical gel preparation of C-82 given daily for 4 weeks. Patient criteria, study design and treatments were as described. After 4 weeks of treatment, skin biopsies were collected from forearms of patients and skin tissue RNA was analyzed using Affymetrix HG-U133A2.0 microarray chips. The clinical trial is registered and publicly available on clinicaltrials.gov #NCT02349009 [39].

**SUPPLEMENTARY LEGENDS:**

**Supplementary figure S1: Validation and visualization of bulk RNA sequencing of 2-day and 5-day βcat$^{istab}$ dermal adipocytes.** A) Validation of Wnt signaling in isolated mature dermal adipocytes RNASeq dataset using Wnt target genes. B) Venn diagram representing total number of DEGs and number of common DEGs in control and 2-, control and 5-, 2- .and 5-day βcat$^{istab}$ adipocytes. C) Heatmap of top 100 differentially expressed genes (DEGs) (sorted by p-value) in 0, 2 and 5-day βcat$^{istab}$ adipocytes obtained from bulk RNASeq D) Heirarchial clustering of all DEGs in control and 2-day and E) control and 5-day βcat$^{istab}$ adipocytes F) DAVID gene ontology (GO) functional annotation clustering based on enrichment (by Fishers exact test) by 2 days and G) 5 days of Wnt activation in mature dermal adipocytes. P-values were calculated with unpaired, two-tailed t-test with Welch's correction when required for unequal variances. *, **, ***, **** is p-value <0.05, 0.01, 0.001 and 0.0001 respectively. A p-value < 0.05 is considered significant.

**Supplementary figure S2: Validation of related lipid metabolism signatures from bulk RNA sequencing of 2-day and 5-day βcat$^{istab}$ dermal adipocytes.** A) FPKM values of control, 2-day and 5-day βcat$^{istab}$ dermal adipocytes for genes associated with adipocyte identity, B) transcription factors involved in *de-novo* lipogenesis, C) genes and enzymes indirectly associated with *de-novo* lipogenesis, D) Fatty acid uptake, E) Lipolysis, F) Fatty acid oxidation and G) Glycolysis from the bulk RNASeq dataset. P-values were calculated with unpaired, two-tailed t-test with Welch's correction when required for unequal variances. *, **, ***, **** is p-value <0.05, 0.01, 0.001 and 0.0001 respectively. A p-value < 0.05 is considered significant.

**Supplementary figure S3: Mining different human sequencing datasets for de novo lipogenesis genes and enzymes.** A) Expression of enzymes in de novo lipogenesis axis and transcription factors from a published human SSc RNASeq dataset GSE130955 sorted by yearly progression of disease. B) Fold change of enzymes in de novo lipogenesis axis, transcription factors/ co-factors and transporters in keloid skin of human patients normalized to adjacent unaffected tissue from published microarray dataset GSE90051. C) Heatmap showing fold change of genes associated with de novo lipogenesis axis post 28 days of treatment with either placebo or topical Wnt inhibitor C-82 lotion. P-values were calculated with unpaired, two-tailed t-test with Welch's correction for two groups. For more than two groups, P-values were calculated using one-way ANOVA with Brown-Forsythe and Welch's ANOVA test and multiple comparisons. *, **, ***, **** is p-value <0.05, 0.01, 0.001 and 0.0001 respectively. A p-value < 0.05 is considered significant.

**Supplementary figure S4: Wnt activation in differentiated adipocytes phenocopies FASN inhibition.** A) Relative quantification of Perilipin1 (PLIN1) and Axin2 mRNA to validate adipocyte identity and Wnt activation in WNT agonist treated cultured differentiated adipocytes (n=3). B) Relative quantity of *de novo* lipogenesis enzyme mRNA *Slc2a4* (GLUT4) and *Acly* with *Hprt* as the housekeeping control. C) Histogram of individual lipid droplet perimeter and D) circularity of differentiated and WNT agonist treated adipocytes (n= fixed 1724 droplets). E) Brightfield, phase contrast and LipidSpot images of undifferentiated, differentiated, 2μM and 20μM WNT agonist, and 200nM TVB3664 (FASN inhibitor) with respective quantification for total lipid droplet area and F) individual lipid droplet area and G) perimeter. H) Lipidomics analysis showing all individual non-esterified (NEFA) species in differentiated, BML-treated and $FASN_i$ treated cultured adipocytes normalized to protein quantity (n=4). I) Lipid product to precursor ratios in all three groups (n=4). For two groups, paired two-tailed Students t-test was employed and for three or more groups, one-way ANOVA with appropriate posthoc test was employed. For the frequency distribution, non-parametric Kruskal-Wallis test was applied. *, **, ***, **** is p-value <0.05, 0.01, 0.001 and 0.0001 respectively. A p-value < 0.05 is considered significant.

**Supplementary figure S5: Downregulation of FASN is accompanied by histomorphometric changes in genetic and chemical model of fibrosis in vivo**. A) Quantification of average individual adipocyte area (microns^2) in p26 control and 5-day βcat$^{istab}$ (n=5) B) Relative quantification of *Acly* and *Acaca* mRNA from DWAT biopsy tissue of p26 control and 5-day βcat$^{istab}$ mouse skin (n=3-4) normalized to *ActB.* C) Histology of 14-day bleomycin (bleo) injected mouse skin with age matched PBS-injected control. D) Immunofluorescence of FASN (red) in PBS- and bleo-injected DWAT and respective quantification of FASN corrected fluorescence (n=3-6). P-values were obtained using unpaired students t-test with Welch's correction for bar and scatter plots. *, **, ***, **** is p-value <0.05, 0.01, 0.001 and 0.0001 respectively. A p-value < 0.05 is considered significant.

**Supplementary figure S6: *Tcf7l2 and Lef1 s*ites on promoter regions of FASN and DNL transcription factors in adipocytes and cell lines with sustained Wnt activation.** A) Integrative Genomics Viewer (IGV) view of *Tcf7l2* binding occupancy in regions ± 3kb from transcription start sites (arrows) of *Srebf1* and *Fasn* in differentiated adipocytes, HCT116 and K562 cell lines in vitro. Scale bar =1kb. B) Predicted transcription factor binding site analysis for human Lef1 motif on *Mlxipl*, *Srebf1* and *Fasn* promoter region containing the Tcf7l2 ChipSeq peak using JASPAR and MEME*.*

**Supplemental figure S7: FASN inhibition during Wnt recovery does not allow recovery of dermal adipocytes.** A) Body weights of p52 control, vehicle treated 10-day reversal $\beta cat^{istab}$ and FASN inhibited 10-day reversal $\beta cat^{istab}$ before and after 10 days of gavage. B) Total thickness of p52 control, 10-day reversal $\beta cat^{istab}$ , 10-day reversal $\beta cat^{istab}$ with vehicle and FASN inhibited 10-day reversal $\beta cat^{istab}$ dorsal skin (n=6-8). C) Histogram of adipocyte perimeter and D) circularity in p52 control, 10-day reversal $\beta cat^{istab}$ and FASN inhibited 10-day reversal $\beta cat^{istab}$ (n=3-6, 394 random cells). E) Lipidomics analysis showing all individual non-esterified (NEFA) species in p52 control, 10-day reversal $\beta cat^{istab}$, 10-day reversal $\beta cat^{istab}$ with vehicle and FASN inhibited 10-day reversal $\beta cat^{istab}$ dorsal skin normalized to tissue weight (n=3). I) Lipid product to precursor ratios in all three groups (n=3)   P-values were determined by 2-way ANOVA with Sidak's test in A), 1-way ANOVA with Tukey's test in B), C) or Fishers LSD test in E) and F) and Kruskal-Wallis test with Dunn's correction in histograms C) and D). *, **, ***, **** is p-value <0.05, 0.01, 0.001 and 0.0001 respectively. A p-value < 0.05 is considered significant.

**Supplementary figure S8: Collagen and highly sulfated proteoglycans in FASN inhibited skin.** A) Principal component analysis (PCA) plot depicting variance of ECM metrics generated by TWOMBLI image analysis algorithm in p52 control, vehicle treated 10-day reversal $\beta cat^{istab}$ and FASN inhibited 10-day reversal $\beta cat^{istab}$. Each point represents one animal. B) TWOMBLI output for coherency or alignment (0 = anisotropy, 1 = isotropy) (n=5-7). C) Alcian blue stained skin (at pH 1.0) to detect highly sulfated proteoglycans and its respective quantification of area occupied by highly sulfated proteoglycans in the upper dermis, lower dermis and DWAT (n=5-8). P-values were obtained by one-way ANOVA with Tukey's multiple comparisons. *, **, ***, **** is p-value <0.05, 0.01, 0.001 and 0.0001 respectively. A p-value < 0.05 is considered significant.

**Supplementary table 1: Targeted lipidomics *in vitro***. Quantification of individual lipid species, segregated into relevant lipid groups and lipid ratios in undifferentiated, differentiated, BML-284 activated and TVB2640 FASN inhibited samples (n=4). P-values were obtained by one-way ANOVA. *, **, ***, **** is p-value <0.05, 0.01, 0.001 and 0.0001 respectively. A p-value < 0.05 is considered significant.

**Supplementary table 2: Targeted lipidomics *in vivo*.** Quantification of individual lipid species, segregated into relevant lipid groups and lipid ratios in p52 control, vehicle treated 10-day reversal $\beta cat^{istab}$ and FASN inhibited 10-day reversal $\beta cat^{istab}$ (n=3). P-values were obtained by one-way ANOVA. *, **, ***, **** is p-value <0.05, 0.01, 0.001 and 0.0001 respectively. A p-value < 0.05 is considered significant.

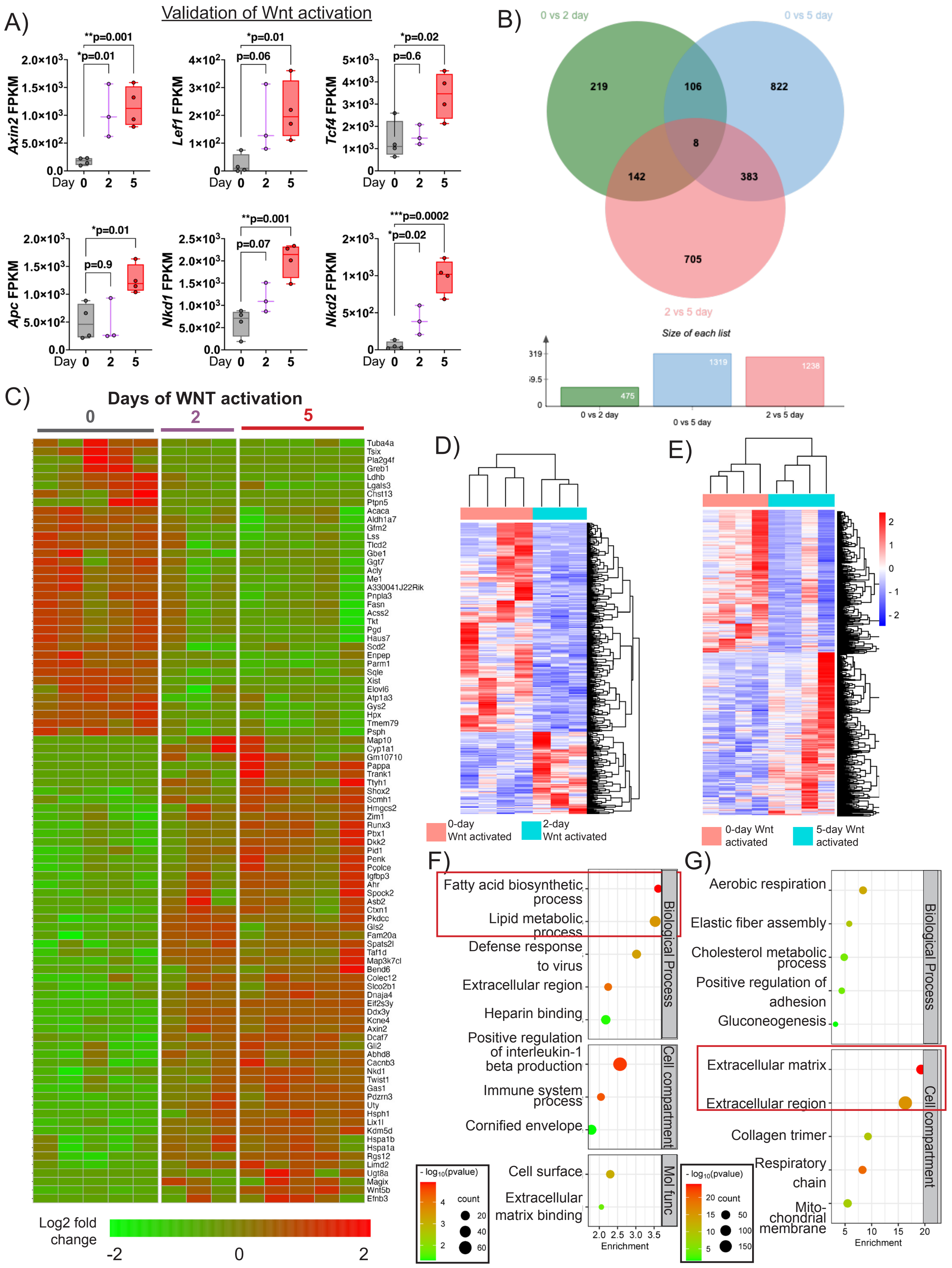
A)
Validation of Wnt activation
Axin2 FPKM
Lef1 FPKM
Tcf4 FPKM
Apc FPKM
Nkd1 FPKM
Nkd2 FPKM
Day 0 2 5
**p=0.001
*p=0.01
p=0.06
*p=0.02
p=0.6
p=0.9
p=0.07
***p=0.0002
B)
0 vs 2 day
0 vs 5 day
2 vs 5 day
219
106
822
8
142
383
705
Size of each list
475
1319
1238
C)
Days of WNT activation
0
2
5
Log2 fold change
-2
0
2
D)
0-day Wnt activated
2-day Wnt activated
E)
0-day Wnt activated
5-day Wnt activated
F)
Fatty acid biosynthetic process
Lipid metabolic process
Defense response to virus
Extracellular region
Heparin binding
Positive regulation of interleukin-1 beta production
Immune system process
Cornified envelope
Cell surface
Extracellular matrix binding
Biological Process
Cell compartment
Mol func
Enrichment
count
G)
Aerobic respiration
Elastic fiber assembly
Cholesterol metabolic process
Positive regulation of adhesion
Gluconeogenesis
Extracellular matrix
Extracellular region
Collagen trimer
Respiratory chain
Mito-chondrial membrane
Biological Process
Cell compartment
Enrichment
count

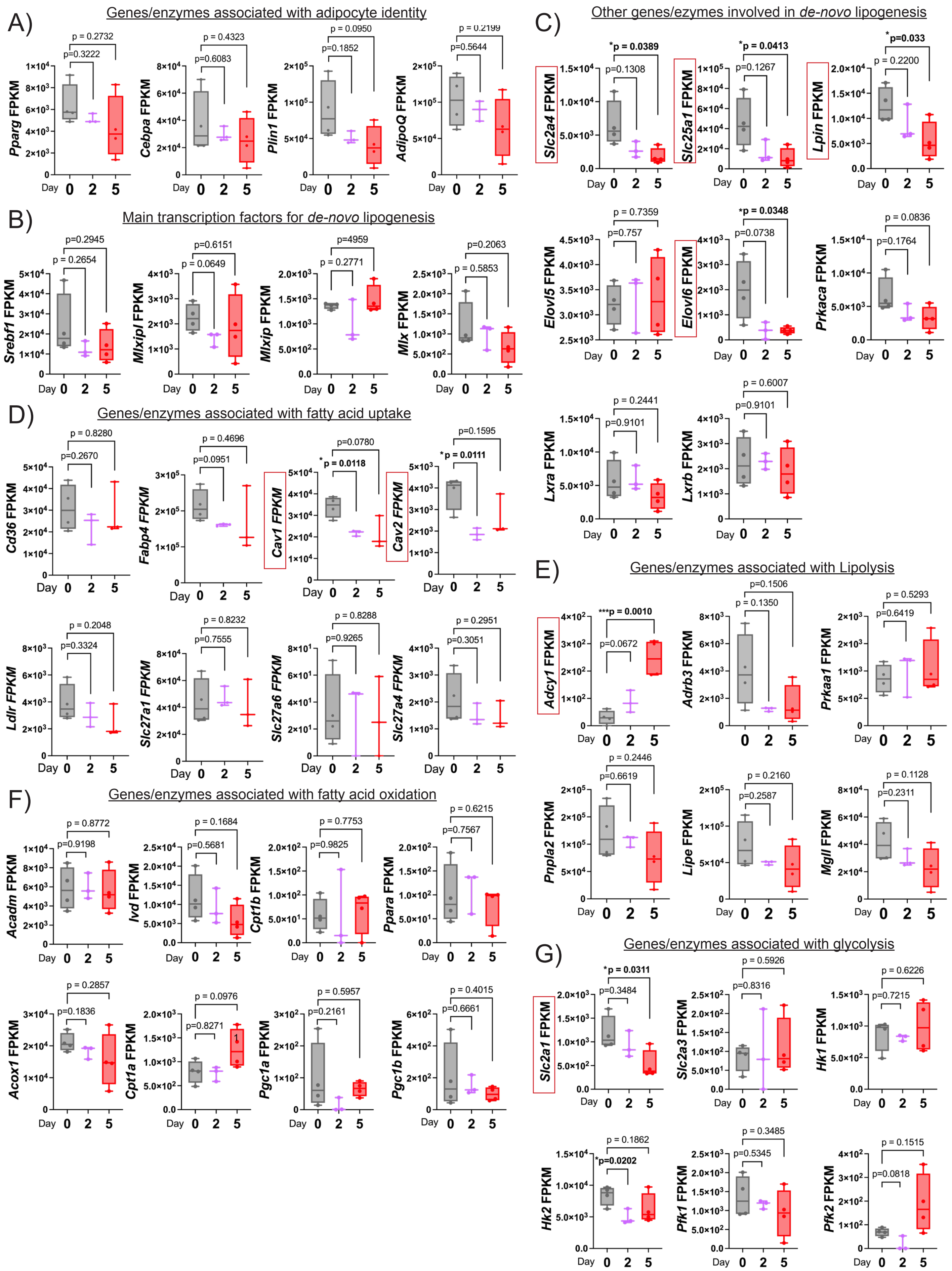
A) Genes/enzymes associated with adipocyte identity
Pparg FPKM
p=0.3222
p = 0.2732
Cebpa FPKM
p=0.6083
p = 0.4323
Plin1 FPKM
p=0.1852
p = 0.0950
AdipoQ FPKM
p=0.5644
p = 0.2199
B) Main transcription factors for de-novo lipogenesis
Srebf1 FPKM
p = 0.2654
p=0.2945
Mlxipl FPKM
p = 0.0649
p=0.6151
Mlxip FPKM
p = 0.2771
p=4959
Mlx FPKM
p = 0.5853
p=0.2063
C) Other genes/ezymes involved in de-novo lipogenesis
Slc2a4 FPKM
p=0.1308
*p = 0.0389
Slc25a1 FPKM
p=0.1267
*p = 0.0413
Lpin FPKM
p = 0.2200
*p=0.033
Elovl5 FPKM
p=0.757
p = 0.7359
Elovl6 FPKM
p=0.0738
*p = 0.0348
Prkaca FPKM
p=0.1764
p = 0.0836
Lxra FPKM
p=0.9101
p = 0.2441
Lxrb FPKM
p=0.9101
p = 0.6007
D) Genes/enzymes associated with fatty acid uptake
Cd36 FPKM
p=0.2670
p = 0.8280
Fabp4 FPKM
p=0.0951
p = 0.4696
Cav1 FPKM
*p = 0.0118
p=0.0780
Cav2 FPKM
*p = 0.0111
p=0.1595
Ldlr FPKM
p=0.3324
p = 0.2048
Slc27a1 FPKM
p=0.7555
p = 0.8232
Slc27a6 FPKM
p=0.9265
p = 0.8288
Slc27a4 FPKM
p=0.3051
p = 0.2951
E) Genes/enzymes associated with Lipolysis
Adcy1 FPKM
p=0.0672
***p = 0.0010
Adrb3 FPKM
p = 0.1350
p=0.1506
Prkaa1 FPKM
p=0.6419
p = 0.5293
Pnpla2 FPKM
p=0.6619
p = 0.2446
Lipe FPKM
p=0.2587
p = 0.2160
Mgll FPKM
p=0.2311
p = 0.1128
F) Genes/enzymes associated with fatty acid oxidation
Acadm FPKM
p=0.9198
p = 0.8772
Ivd FPKM
p=0.5681
p = 0.1684
Cpt1b FPKM
p=0.9825
p = 0.7753
Ppara FPKM
p=0.7567
p = 0.6215
Acox1 FPKM
p=0.1836
p = 0.2857
Cpt1a FPKM
p=0.8271
p = 0.0976
Pgc1a FPKM
p=0.2161
p = 0.5957
Pgc1b FPKM
p=0.6661
p = 0.4015
G) Genes/enzymes associated with glycolysis
Slc2a1 FPKM
p=0.3484
*p = 0.0311
Slc2a3 FPKM
p=0.8316
p = 0.5926
Hk1 FPKM
p=0.7215
p = 0.6226
Hk2 FPKM
*p=0.0202
p = 0.1862
Pfk1 FPKM
p=0.5345
p = 0.3485
Pfk2 FPKM
p=0.0818
p = 0.1515
Day 0 2 5

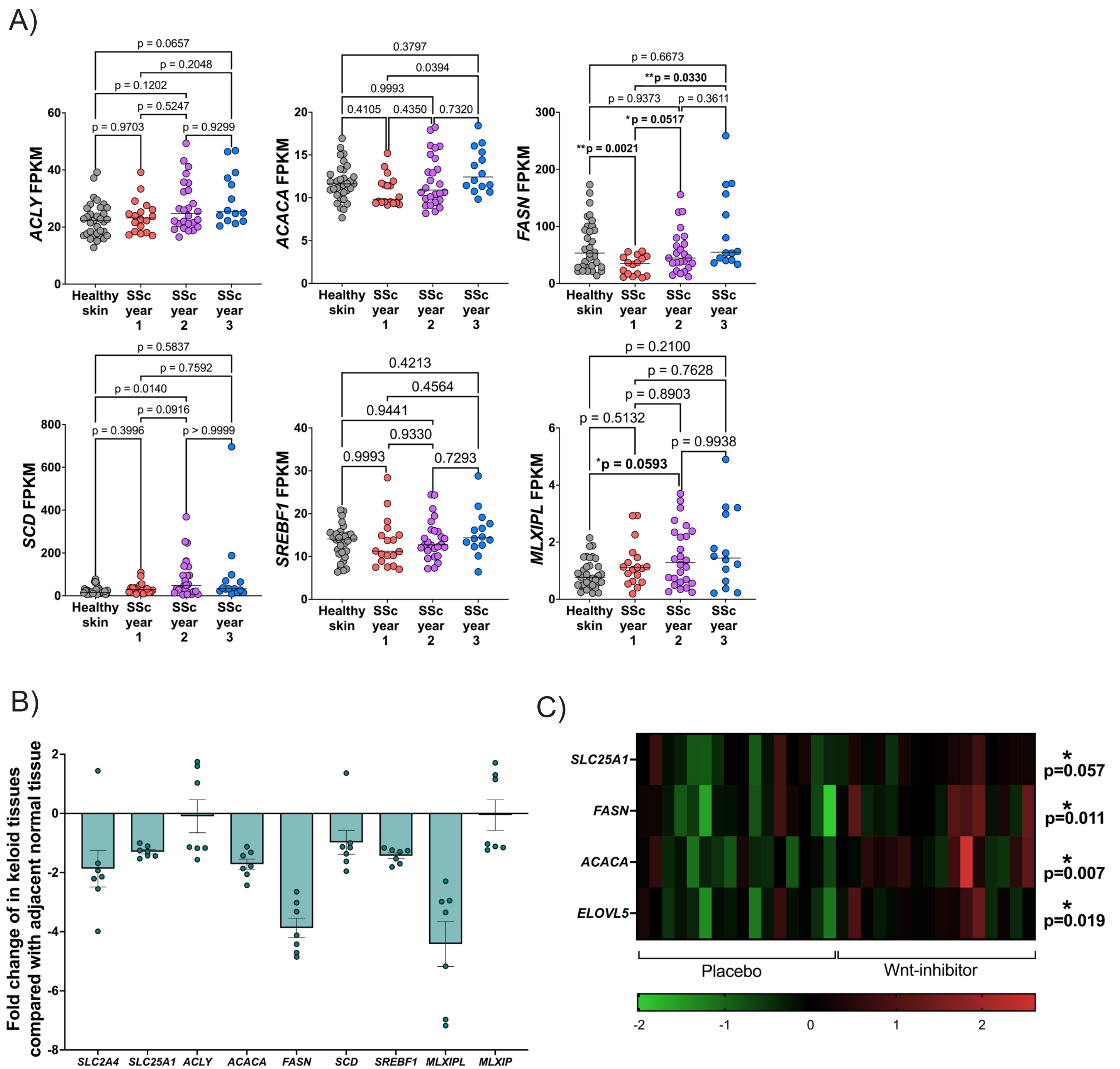
A)
ACLY FPKM
ACACA FPKM
FASN FPKM
SCD FPKM
SREBF1 FPKM
MLXIPL FPKM
Healthy skin
SSc year 1
SSc year 2
SSc year 3
p = 0.0657
p = 0.2048
p = 0.1202
p = 0.5247
p = 0.9703
p = 0.9299
0.3797
0.0394
0.9993
0.4105
0.4350
0.7320
p = 0.6673
**p = 0.0330
p = 0.9373
p = 0.3611
*p = 0.0517
**p = 0.0021
p = 0.5837
p = 0.7592
p = 0.0140
p = 0.0916
p = 0.3996
p > 0.9999
0.4213
0.4564
0.9441
0.9330
0.9993
0.7293
p = 0.2100
p = 0.7628
p = 0.8903
p = 0.5132
p = 0.9938
*p = 0.0593
B)
Fold change of in keloid tissues compared with adjacent normal tissue
SLC2A4
SLC25A1
ACLY
ACACA
FASN
SCD
SREBF1
MLXIPL
MLXIP
C)
SLC25A1
FASN
ACACA
ELOVL5
* p=0.057
* p=0.011
* p=0.007
* p=0.019
Placebo
Wnt-inhibitor
-2
-1
0
1
2

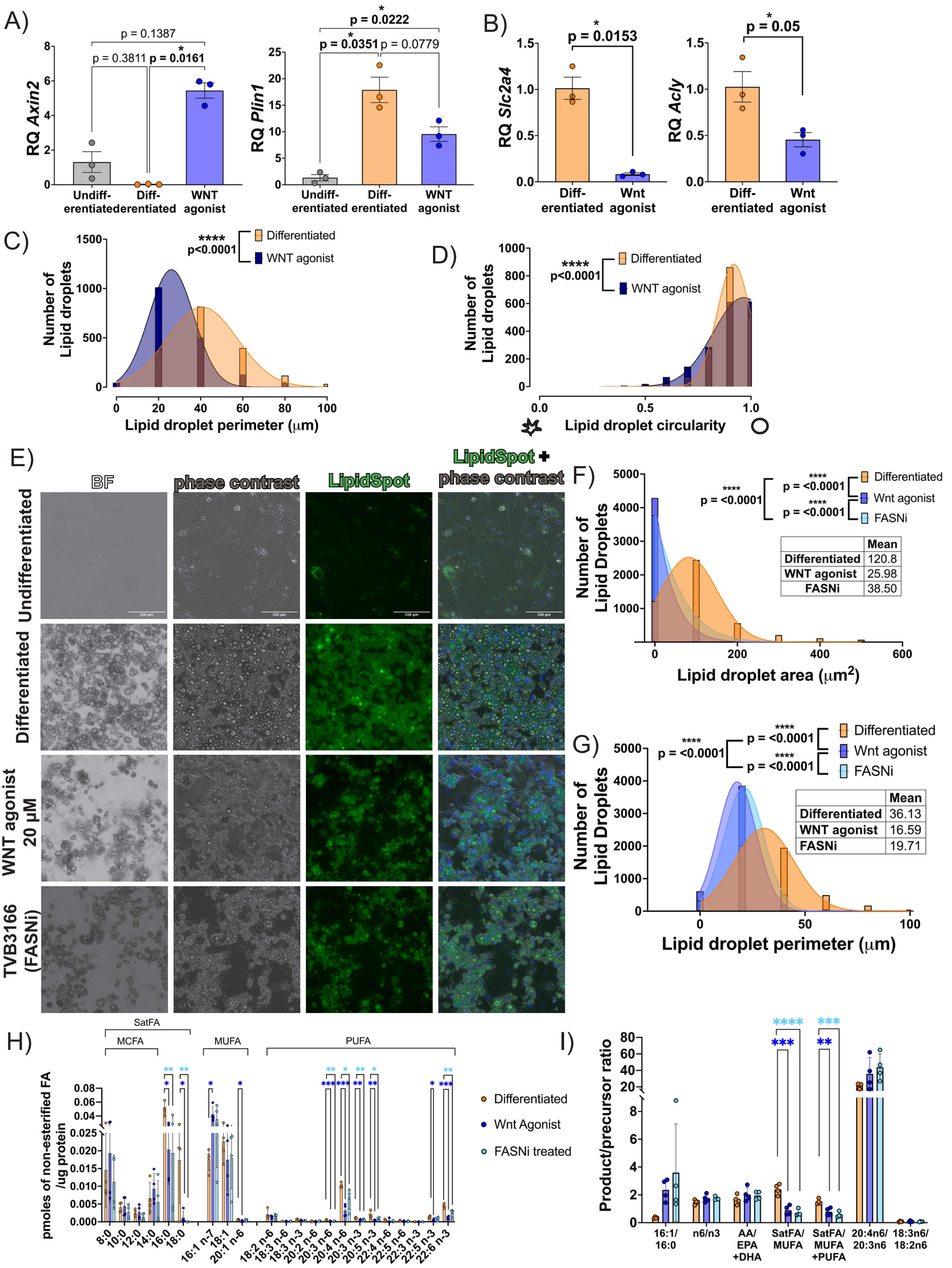

A)
RQ Axin2
p = 0.1387
p = 0.3811
p = 0.0161
Undiff-erentiated
Diff-erentiated
WNT agonist
RQ Plin1
p = 0.0222
p = 0.0351
p = 0.0779
B)
RQ Slc2a4
p = 0.0153
Diff-erentiated
Wnt agonist
RQ Acly
p = 0.05
C)
Number of Lipid droplets
**** p<0.0001
Differentiated
WNT agonist
Lipid droplet perimeter (μm)
D)
Lipid droplet circularity
E)
BF
phase contrast
LipidSpot
LipidSpot + phase contrast
Undifferentiated
Differentiated
WNT agonist 20 μM
TVB3166 (FASNi)
F)
Number of Lipid Droplets
p = <0.0001
Wnt agonist
FASNi
Lipid droplet area (μm2)
Mean
Differentiated 120.8
WNT agonist 25.98
FASNi 38.50
G)
Lipid droplet perimeter (μm)
Differentiated 36.13
WNT agonist 16.59
FASNi 19.71
H)
SatFA
MCFA
MUFA
PUFA
pmoles of non-esterified FA /ug protein
8:0
10:0
12:0
14:0
16:0
18:0
16:1 n-7
18:1
20:1 n-6
18:2 n-6
18:3 n-6
18:3 n-3
20:2 n-6
20:3 n-6
20:4 n-6
20:3 n-3
20:5 n-3
22:4 n-6
22:5 n-6
22:3 n-3
22:5 n-3
22:6 n-3
Differentiated
Wnt Agonist
FASNi treated
I)
Product/precursor ratio
16:1/16:0
n6/n3
AA/EPA+DHA
SatFA/MUFA
SatFA/MUFA+PUFA
20:4n6/20:3n6
18:3n6/18:2n6

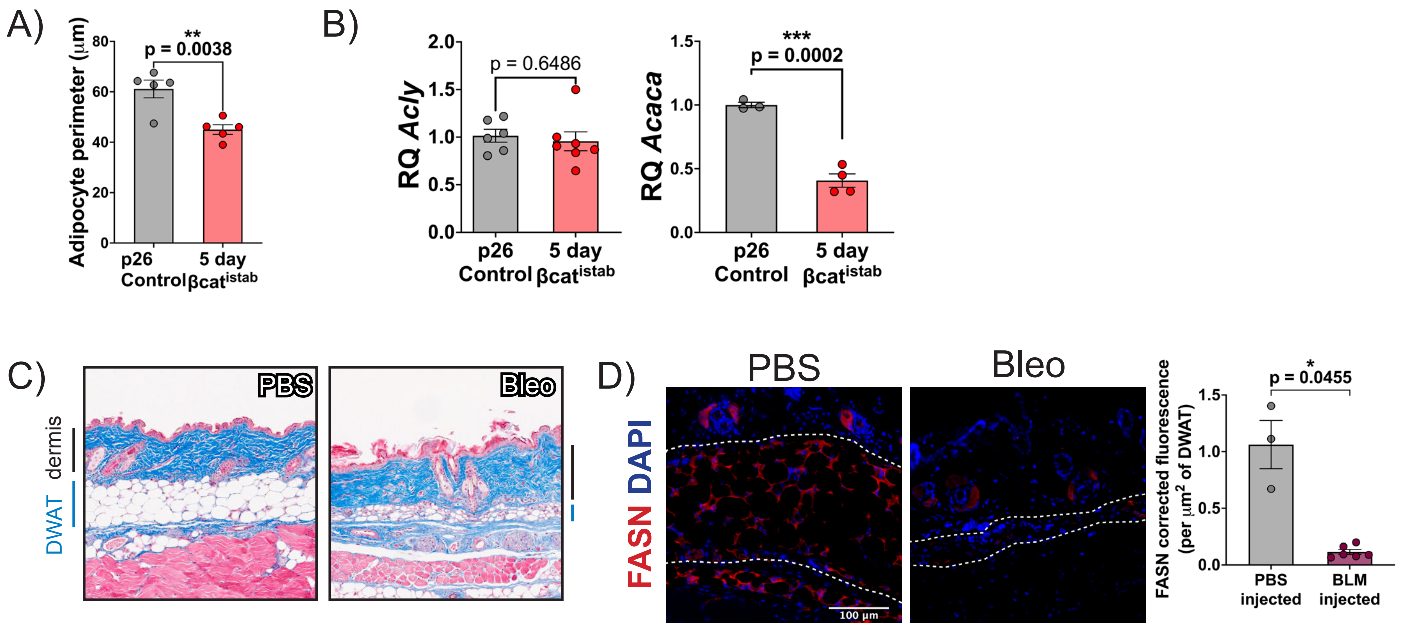

A)
Adipocyte perimeter (μm)
p = 0.0038
**
p26 Control
5 day βcat istab
B)
RQ Acly
p = 0.6486
RQ Acaca
***
p = 0.0002
p26 Control
5 day βcat istab
C)
PBS
Bleo
dermis
DWAT
D)
PBS
Bleo
FASN DAPI
100 μm
FASN corrected fluorescence (per μm2 of DWAT)
*
p = 0.0455
PBS injected
BLM injected

A)

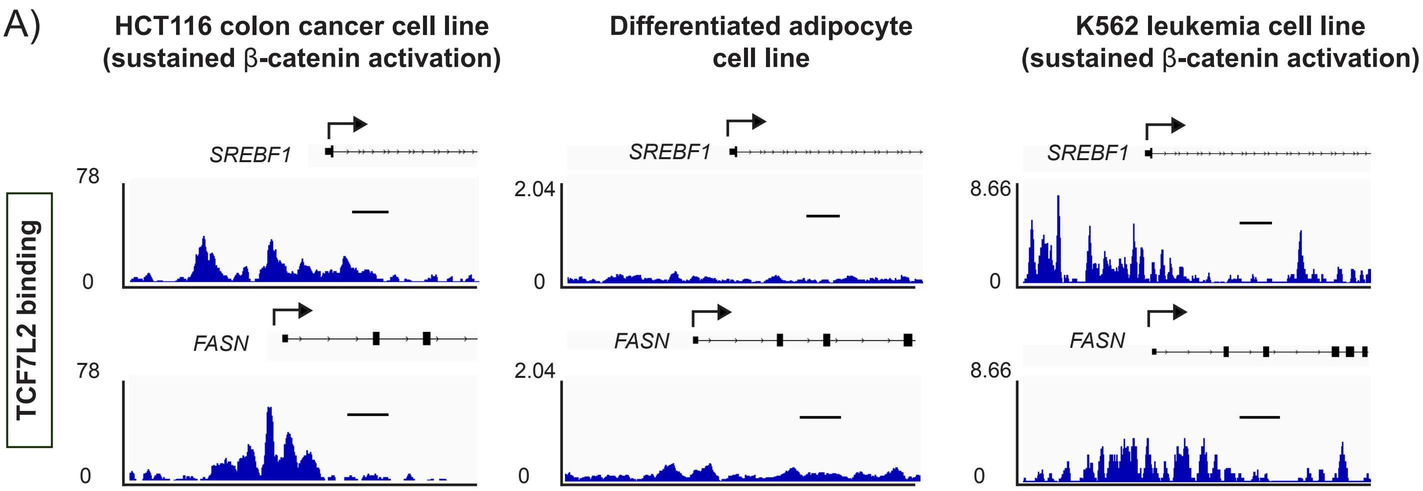


B)

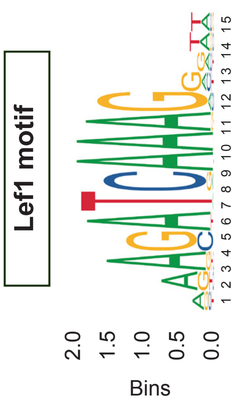


| motif_id | gene_ name | start | stop | strand | score | p-value | q-value | matched_sequence | |
|---|---|---|---|---|---|---|---|---|---|
| **MA076 8.1 LEF1** | *Mlxipl* range= chr7 | 73037615 | 73037629 | + | 9.69355 | 4.56E-05 | 0.209 | AAAGATCAAGGCCGG | +++++++++ + <br> CAGAGAAAAGATCAAGGCCGGCCGAT |
| **MA076 8.1 LEF1** | *Srebf1* range= chr17 | 17741601 | 17741615 | - | 9.03226 | 5.91E-05 | 0.399 | GAGGAACAAAGGGAG | ++++++++ ++ + <br> CCAGCTCCCTTTGTTCCTCTCCC |
| **MA076 8.1 LEF1** | *Fasn* range= chr17 | 80054793 | 80054807 | + | 8.53226 | 7.18E-05 | 0.389 | GGCCATCAAAGAAAT | +++++++ +++ <br> TAAGGGCCATCAAAGAAATGGAGA |

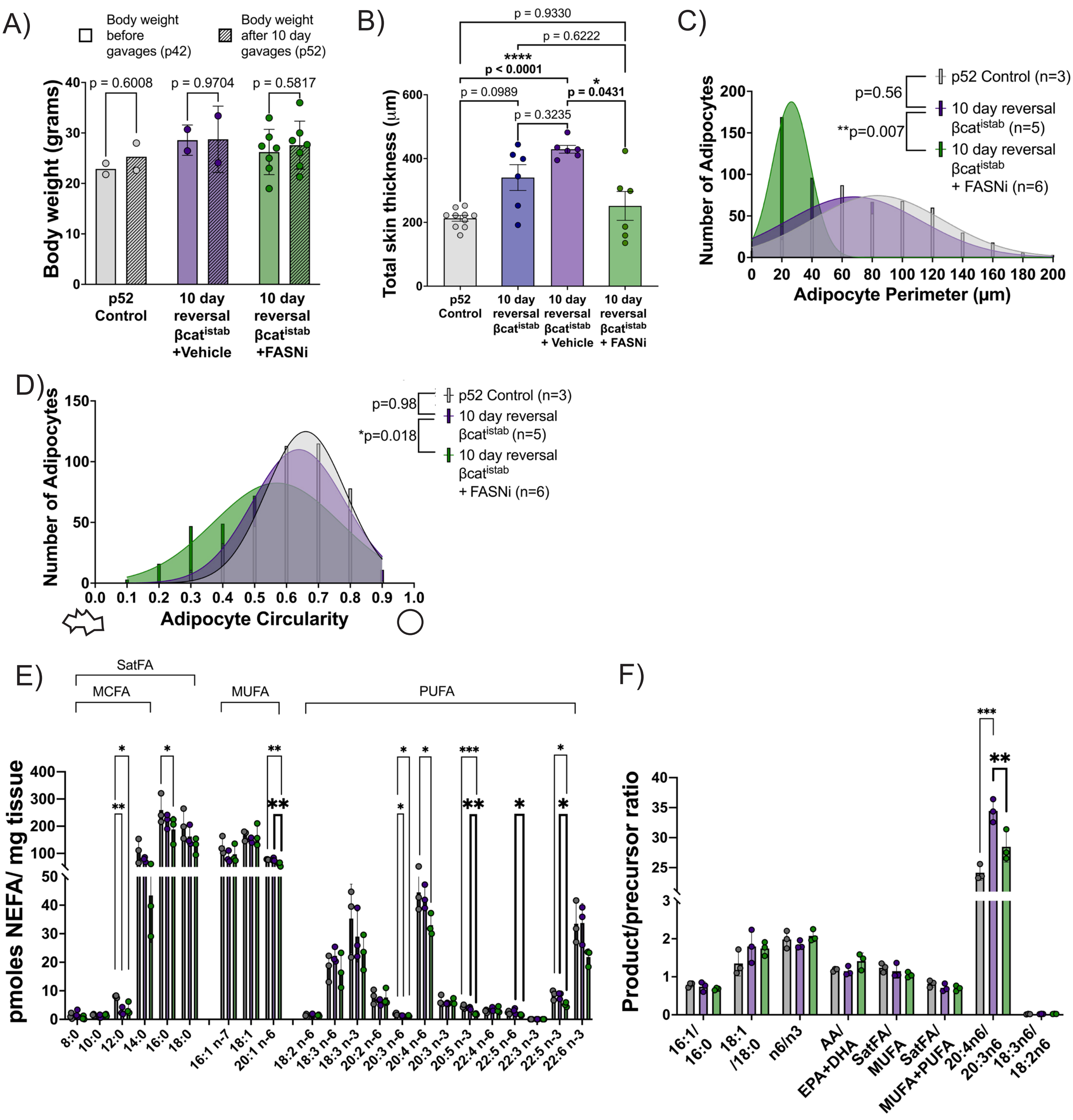
A)
Body weight before gavages (p42)
Body weight after 10 day gavages (p52)
p = 0.6008
p = 0.9704
p = 0.5817
Body weight (grams)
p52 Control
10 day reversal βcat istab +Vehicle
10 day reversal βcat istab +FASNi
B)
p = 0.9330
p = 0.6222
****
p < 0.0001
p = 0.0989
*
p = 0.0431
p = 0.3235
Total skin thickness (μm)
p52 Control
10 day reversal βcat istab
10 day reversal βcat istab + Vehicle
10 day reversal βcat istab + FASNi
C)
p52 Control (n=3)
p=0.56
10 day reversal βcat istab (n=5)
**p=0.007
10 day reversal βcat istab + FASNi (n=6)
Number of Adipocytes
Adipocyte Perimeter (μm)
D)
p52 Control (n=3)
p=0.98
10 day reversal βcat istab (n=5)
*p=0.018
10 day reversal βcat istab + FASNi (n=6)
Number of Adipocytes
Adipocyte Circularity
E)
SatFA
MCFA
MUFA
PUFA
pmoles NEFA/ mg tissue
8:0
10:0
12:0
14:0
16:0
18:0
16:1 n-7
18:1
20:1 n-6
18:2 n-6
18:3 n-6
18:3 n-3
20:2 n-6
20:3 n-6
20:4 n-6
20:3 n-3
20:5 n-3
22:4 n-6
22:5 n-6
22:3 n-3
22:5 n-3
22:6 n-3
F)
Product/precursor ratio
16:1/ 16:0
18:1 /18:0
n6/n3
AA/ EPA+DHA
SatFA/ MUFA
SatFA/ MUFA+PUFA
20:4n6/ 20:3n6
18:3n6/ 18:2n6

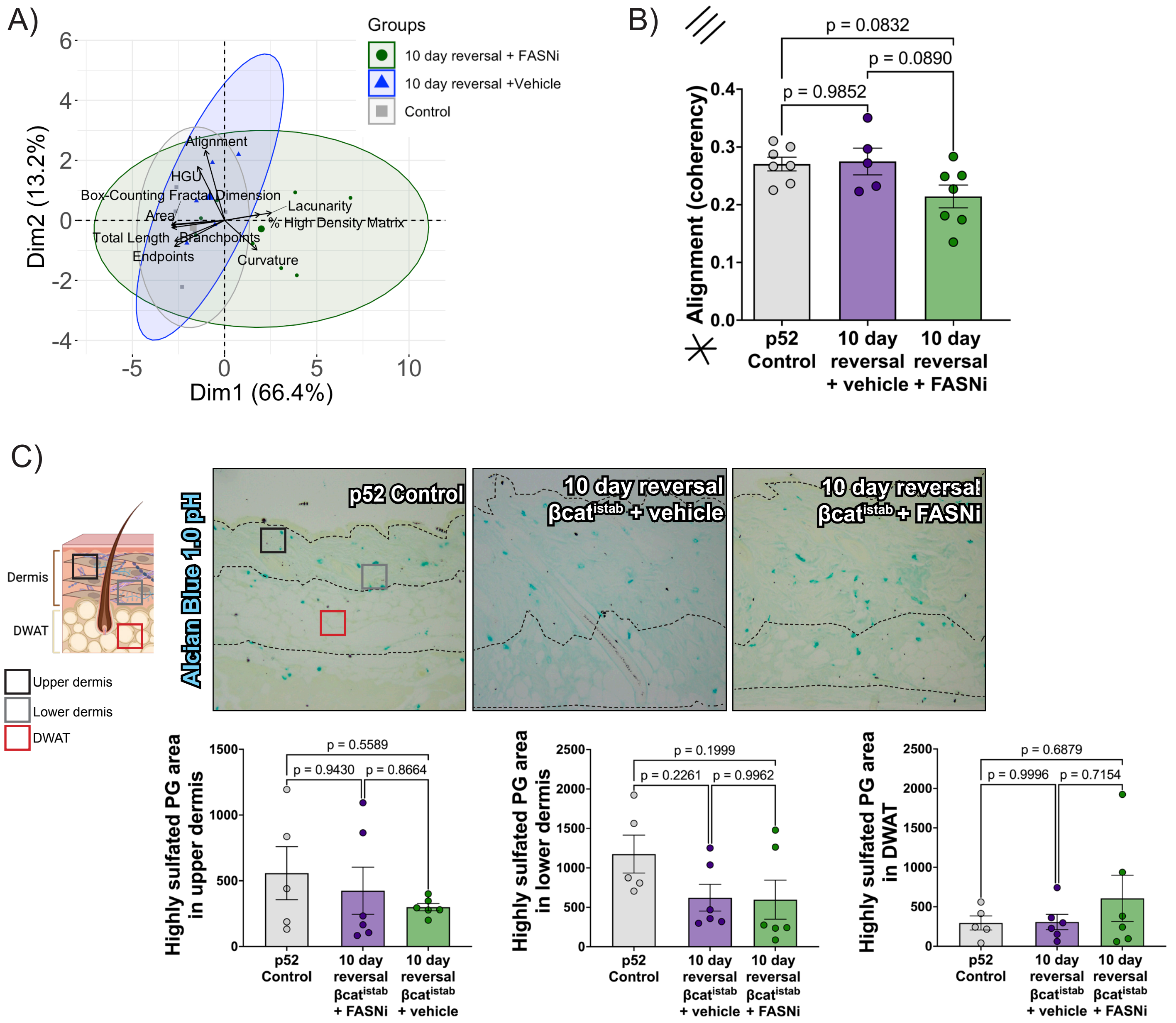
A)
Groups
10 day reversal + FASNi
10 day reversal +Vehicle
Control
Dim2 (13.2%)
Dim1 (66.4%)
Alignment
HGU
Box-Counting Fractal Dimension
Lacunarity
Area
% High Density Matrix
Total Length
Branchpoints
Endpoints
Curvature
B)
p = 0.0832
p = 0.0890
p = 0.9852
Alignment (coherency)
p52 Control
10 day reversal + vehicle
10 day reversal + FASNi
C)
Dermis
DWAT
Upper dermis
Lower dermis
DWAT
Alcian Blue 1.0 pH
p52 Control
10 day reversal βcatistab + vehicle
10 day reversal βcatistab + FASNi
Highly sulfated PG area in upper dermis
p = 0.5589
p = 0.9430
p = 0.8664
p52 Control
10 day reversal βcatistab + FASNi
10 day reversal βcatistab + vehicle
Highly sulfated PG area in lower dermis
p = 0.1999
p = 0.2261
p = 0.9962
p52 Control
10 day reversal βcatistab + vehicle
10 day reversal βcatistab + FASNi
Highly sulfated PG area in DWAT
p = 0.6879
p = 0.9996
p = 0.7154
p52 Control
10 day reversal βcatistab + vehicle
10 day reversal βcatistab + FASNi

**Table 1.1 Targeted lipidomics analysis on the non-esterified fatty acid (NEFA) lipid fraction in undifferentiated, differentiated, wnt activated and FASN inhibited adipocyte cell culture samples**

| | Undifferentiated | | Differentiated | | | | Wnt agonist | | | | FASNi treated | | | |
|---|---|---|---|---|---|---|---|---|---|---|---|---|---|---|
| ID | Mean | SD | Mean | SD | | Fold change vs Undifferentiated | Mean | SD | | Fold change vs Differentiated | Mean | SD | | Fold change vs Differentiated |
| 10:0 | 12.61 | 0.36 | 16.34 | 4.06 | | 1.30 | 13.80 | 3.47 | | 0.84 | 10.26 | 1.15 | # | 0.63 |
| 12:0 | 7.32 | 1.67 | 9.95 | 3.77 | | 1.36 | 8.34 | 3.40 | | 0.84 | 5.03 | 1.20 | | 0.51 |
| 14:0 | 90.12 | 11.73 | 23.59 | 4.07 | **** | 0.26 | 32.24 | 9.52 | **** | 1.37 | 25.41 | 22.44 | **** | 1.08 |
| 14:1 | 4.62 | 1.47 | 4.24 | 2.30 | | 0.92 | 27.87 | 14.16 | * # | 6.58 | 23.05 | 10.42 | # | 5.44 |
| 16:0 | 1471.55 | 117.79 | 219.55 | 80.68 | **** | 0.15 | 68.43 | 12.49 | **** | 0.31 | 92.75 | 80.98 | **** | 0.42 |
| 16:1 n-7 | 43.73 | 40.88 | 74.04 | 22.70 | | 1.69 | 164.32 | 64.13 | | 2.22 | 167.60 | 106.80 | | 2.26 |
| 18:0 | 607.07 | 49.02 | 90.64 | 67.23 | **** | 0.15 | 3.29 | 4.88 | **** | 0.04 | 0.00 | 0.00 | **** # | 0.00 |
| 18:1 | 178.82 | 77.26 | 92.94 | 27.14 | | 0.52 | 59.37 | 19.01 | * | 0.64 | 83.46 | 52.36 | | 0.90 |
| 18:2 n-6 | 15.54 | 1.74 | 9.73 | 8.40 | | 0.63 | 5.28 | 2.64 | * | 0.54 | 7.18 | 3.79 | | 0.74 |
| 18:3 n-6 | 0.60 | 0.28 | 0.55 | 0.24 | | 0.91 | 0.40 | 0.14 | | 0.73 | 0.48 | 0.14 | | 0.87 |
| 18:3 n-3 | 2.03 | 0.31 | 1.80 | 1.29 | | 0.89 | 1.64 | 0.90 | | 0.91 | 1.50 | 0.73 | | 0.83 |
| 20:1 n-6 | 9.57 | 7.26 | 2.32 | 0.76 | | 0.24 | 0.74 | 1.14 | * | 0.32 | 1.94 | 1.13 | | 0.83 |
| 20:2 n-6 | 2.93 | 1.45 | 0.44 | 0.53 | ** | 0.15 | 0.12 | 0.25 | ** | 0.28 | 0.65 | 0.40 | ** | 1.50 |
| 20:3 n-6 | 2.70 | 1.85 | 2.20 | 1.09 | | 0.81 | 0.35 | 0.15 | | 0.16 | 0.80 | 0.63 | | 0.37 |
| 20:4 n-6 | 24.41 | 8.55 | 45.61 | 18.88 | | 1.87 | 10.41 | 1.22 | ## | 0.23 | 29.03 | 12.29 | | 0.64 |
| 20:3 n-3 | 4.51 | 1.69 | 4.87 | 2.36 | | 1.08 | 1.39 | 0.32 | # | 0.28 | 2.37 | 0.71 | | 0.49 |
| 20:5 n-3 | 1.43 | 1.20 | 9.95 | 8.10 | | 6.93 | 1.46 | 0.37 | | 0.15 | 4.22 | 3.21 | | 0.42 |
| 22:4 n-6 | 2.48 | 1.48 | 2.05 | 1.38 | | 0.83 | 0.53 | 1.06 | | 0.26 | 0.70 | 1.33 | | 0.34 |
| 22:5 n-6 | 1.05 | 0.32 | 1.22 | 0.61 | | 1.16 | 0.31 | 0.41 | | 0.25 | 0.71 | 0.35 | | 0.58 |
| 22:5 n-3 | 6.98 | 2.64 | 5.47 | 1.38 | | 0.78 | 1.77 | 0.33 | ** # | 0.32 | 3.36 | 1.31 | * | 0.61 |
| 22:6 n-3 | 15.71 | 6.86 | 20.17 | 9.29 | | 1.28 | 3.94 | 0.97 | # | 0.20 | 11.44 | 5.44 | | 0.57 |
| MCFA | 134.66 | 10.06 | 101.27 | 7.69 | | 0.75 | 112.36 | 32.14 | | 1.11 | 84.12 | 25.51 | * | 0.83 |
| SatFA | 2213.28 | 175.25 | 411.46 | 139.98 | **** | 0.19 | 184.08 | 46.57 | **** | 0.45 | 176.87 | 103.46 | **** | 0.43 |
| MUFA | 232.12 | 124.60 | 169.30 | 49.70 | | 0.73 | 224.42 | 75.07 | | 1.33 | 253.00 | 157.75 | | 1.49 |
| PUFA | 80.84 | 25.58 | 104.05 | 48.09 | | 1.29 | 27.60 | 5.07 | # | 0.27 | 62.45 | 29.17 | | 0.60 |
| LC-PUFA | 62.67 | 24.52 | 91.97 | 41.24 | | 1.47 | 20.27 | 2.17 | # | 0.22 | 53.30 | 24.59 | | 0.58 |
| n-6 PUFA | 49.70 | 13.29 | 61.79 | 28.68 | | 1.24 | 17.40 | 2.89 | # | 0.28 | 39.55 | 18.21 | | 0.64 |
| n-3 PUFA | 31.15 | 12.70 | 42.26 | 20.08 | | 1.36 | 10.20 | 2.41 | # | 0.24 | 22.90 | 11.01 | | 0.54 |
| n6:n3 | 1.68 | 0.34 | 1.48 | 0.17 | | 0.88 | 1.75 | 0.28 | | 1.18 | 1.74 | 0.12 | | 1.18 |
| AA:EPA+DHA | 1.49 | 0.28 | 1.65 | 0.34 | | 1.11 | 2.02 | 0.51 | | 1.22 | 1.96 | 0.29 | | 1.19 |
| SatFA:MUFA | 11.59 | 5.26 | 2.40 | 0.38 | ** | 0.21 | 0.89 | 0.34 | **** | 0.37 | 0.71 | 0.23 | **** | 0.29 |
| SatFA:MUFA+PUFA | 8.15 | 3.09 | 1.51 | 0.18 | *** | 0.19 | 0.79 | 0.28 | *** | 0.52 | 0.56 | 0.20 | **** | 0.37 |
| 18:1/18:0 | 0.29 | 0.10 | 0.86 | 0.10 | | 2.99 | 33.12 | 23.11 | * # | 38.32 | N/A | N/A | | N/A |
| 16:1/16:0 | 0.03 | 0.03 | 0.35 | 0.06 | | 12.38 | 2.36 | 0.79 | | 6.73 | 3.60 | 3.50 | | 10.27 |
| 20:4n-6/20:3n-6 | 11.47 | 6.67 | 21.44 | 2.91 | | 1.87 | 35.97 | 19.55 | | 1.68 | 43.86 | 16.18 | * | 2.05 |
| 18:3n-6/18:2n-6 | 0.04 | 0.02 | 0.07 | 0.03 | | 1.93 | 0.09 | 0.04 | | 1.15 | 0.07 | 0.02 | | 0.98 |
| 14:1/14:0 | 0.05 | 0.02 | 0.17 | 0.07 | | 3.31 | 0.87 | 0.36 | * | 5.13 | 3.42 | 4.83 | * # | 20.09 |
| Total FA | 2530.86 | 318.79 | 689.05 | 232.01 | **** | 0.27 | 463.97 | 111.24 | **** | 0.67 | 515.36 | 291.68 | **** | 0.75 |

*** Significant with respect to Undifferentiated, one-way Anova**
**# Significant with respect to Differentiated, one-way Anova**
**$ Significant with respect to Wnt agonist, one-way Anova**
**All values in pmoles/mg tissue**

**Table 1.2 Targeted lipidomics analysis on the total fatty acid (non-esterified+esterified) lipid fraction in undifferentiated, differentiated, wnt activated and FASN inhibited adipocyte cell culture samples**

| | Undifferentiated | | Differentiated | | | | Wnt agonist | | | | FASNi treated | | | |
|---|---|---|---|---|---|---|---|---|---|---|---|---|---|---|
| ID | Mean | SD | Mean | SD | | Fold change vs Undifferentiated | Mean | SD | | Fold change vs Differentiated | Mean | SD | | Fold change vs Differentiated |
| 10:0 | 0 | 0 | 236 | 178 | * | N/A | 35 | 31 | # | 0.15 | 0 | 0 | # | N/A |
| 12:0 | 18 | 19 | 604 | 484 | * | 34.20 | 96 | 76 | | 0.16 | 2 | 4 | # | 0.00 |
| 14:0 | 111 | 36 | 8429 | 3193 | *** | 75.78 | 2493 | 1588 | ## | 0.30 | 569 | 363 | ### | 0.07 |
| 14:1 | 8 | 2 | 9199 | 5425 | ** | 1190.05 | 1724 | 1084 | # | 0.19 | 653 | 276 | ## | 0.07 |
| 16:0 | 1332 | 452 | 12218 | 4958 | *** | 9.17 | 3592 | 1961 | ## | 0.29 | 823 | 662 | ### | 0.07 |
| 16:1 n-7 | 119 | 63 | 18261 | 8293 | *** | 152.96 | 5171 | 2365 | ## | 0.28 | 3531 | 969 | ## | 0.19 |
| 18:0 | 1026 | 261 | 2468 | 1827 | | 2.41 | 155 | 219 | # | 0.06 | 0 | 0 | # | N/A |
| 18:1 | 374 | 248 | 10943 | 8585 | * | 29.25 | 2750 | 1627 | | 0.25 | 1360 | 462 | # | 0.12 |
| 18:2 n-6 | 28 | 8 | 534 | 228 | *** | 19.33 | 155 | 82 | ## | 0.29 | 71 | 21 | ### | 0.13 |
| 18:3 n-6 | 2 | 0 | 60 | 30 | ** | 28.43 | 16 | 10 | # | 0.27 | 6 | 1 | ## | 0.10 |
| 18:3 n-3 | 6 | 3 | 56 | 33 | ** | 9.45 | 12 | 8 | # | 0.21 | 8 | 3 | ## | 0.15 |
| 20:1 n-6 | 9 | 9 | 485 | 257 | ** | 55.25 | 85 | 61 | ## | 0.17 | 54 | 27 | ## | 0.11 |
| 20:2 n-6 | 3 | 3 | 29 | 11 | *** | 8.69 | 6 | 3 | ### | 0.22 | 4 | 2 | ### | 0.14 |
| 20:3 n-6 | 2 | 2 | 30 | 17 | ** | 15.19 | 4 | 2 | ## | 0.12 | 2 | 2 | ## | 0.06 |
| 20:4 n-6 | 39 | 16 | 459 | 254 | ** | 11.88 | 54 | 20 | ## | 0.12 | 40 | 20 | ## | 0.09 |
| 20:3 n-3 | 4 | 3 | 62 | 36 | ** | 13.94 | 14 | 7 | # | 0.23 | 6 | 3 | ## | 0.09 |
| 20:5 n-3 | 4 | 2 | 80 | 52 | ** | 20.31 | 10 | 6 | # | 0.13 | 8 | 4 | ## | 0.10 |
| 22:4 n-6 | 3 | 2 | 31 | 13 | *** | 11.17 | 7 | 5 | ## | 0.24 | 1 | 1 | ### | 0.03 |
| 22:5 n-6 | 2 | 2 | 18 | 9 | ** | 7.77 | 3 | 2 | ## | 0.18 | 2 | 1 | ## | 0.12 |
| 22:5 n-3 | 12 | 1 | 86 | 45 | ** | 7.34 | 17 | 6 | ## | 0.20 | 9 | 5 | ## | 0.10 |
| 22:6 n-3 | 17 | 9 | 247 | 143 | ** | 14.96 | 43 | 23 | ## | 0.18 | 21 | 11 | ## | 0.09 |
| MCFA | 129 | 50 | 9270 | 3837 | *** | 71.93 | 2634 | 1679 | ## | 0.28 | 575 | 360 | ### | 0.06 |
| SatFA | 2487 | 722 | 23956 | 10412 | *** | 9.63 | 6380 | 3777 | ## | 0.27 | 1398 | 1012 | ### | 0.06 |
| MUFA | 502 | 314 | 29689 | 16284 | ** | 59.11 | 8006 | 4051 | # | 0.27 | 4945 | 1454 | ## | 0.17 |
| PUFA | 124 | 40 | 1694 | 838 | ** | 13.65 | 345 | 171 | ## | 0.20 | 181 | 61 | ## | 0.11 |
| LC-PUFA | 88 | 31 | 1045 | 559 | ** | 11.81 | 162 | 72 | ## | 0.16 | 95 | 45 | ## | 0.09 |
| n-6 PUFA | 79 | 27 | 1161 | 542 | *** | 14.74 | 246 | 123 | ## | 0.21 | 126 | 44 | ## | 0.11 |
| n-3 PUFA | 45 | 13 | 533 | 303 | ** | 11.77 | 99 | 48 | ## | 0.19 | 54 | 18 | ## | 0.10 |
| n6:n3 | 1.72 | 0.11 | 2.33 | 0.38 | ** | 1.35 | 2.43 | 0.14 | ** | 1.04 | 2.33 | 0.13 | * | 1.00 |
| AA:EPA+DHA | 1.93 | 0.36 | 1.48 | 0.25 | | 0.77 | 1.17 | 0.41 | * | 0.79 | 1.38 | 0.11 | | 0.93 |
| SatFA:MUFA | 5.84 | 1.97 | 0.83 | 0.13 | **** | 0.14 | 0.76 | 0.11 | **** | 0.91 | 0.25 | 0.13 | **** | 0.30 |
| SatFA:MUFA+PUFA | 4.40 | 1.18 | 0.79 | 0.11 | **** | 0.18 | 0.73 | 0.11 | **** | 0.93 | 0.24 | 0.13 | **** | 0.31 |
| 18:1/18:0 | 0.34 | 0.14 | 5.14 | 2.59 | | 15.20 | 36.53 | 26.95 | * # | 7.11 | N/A | N/A | | N/A |
| 16:1/16:0 | 0.09 | 0.02 | 1.49 | 0.14 | | 17.41 | 1.50 | 0.18 | | 1.01 | 6.84 | 4.48 | ** # $ | 4.60 |
| 20:4n-6/20:3n-6 | 51.38 | 62.21 | 15.76 | 1.77 | | 0.31 | 34.41 | 41.47 | | 2.18 | 17.95 | 1.12 | | 1.14 |
| 18:3n-6/18:2n-6 | 0.08 | 0.02 | 0.11 | 0.02 | | 1.35 | 0.10 | 0.03 | | 0.89 | 0.09 | 0.03 | | 0.84 |
| 14:1/14:0 | 0.07 | 0.03 | 1.00 | 0.36 | * | 13.79 | 0.68 | 0.15 | | 0.68 | 1.42 | 0.68 | ** | 1.42 |
| Total FA | 3113 | 1048 | 55340 | 27034 | ** | 17.78 | 14731 | 7953 | ## | 0.27 | 6523 | 2517 | ## | 0.12 |

*** Significant with respect to Undifferentiated, one-way Anova**
**# Significant with respect to Differentiated, one-way Anova**
**$ Significant with respect to Wnt agonist, one-way Anova**
**All values in pmoles/mg tissue**

**Table 2.1** Targeted lipidomics analysis on the non-esterified fatty acid (NEFA) lipid fraction from dorsal skin of p52 controls, vehicle-treated 10-day reversal $\beta cat^{istab}$ and FASN inhibited 10-day reversal $\beta cat^{istab}$

| | **p52 Controls** | | 10 day reversal $\beta cat^{istab}$ **+ vehicle** | | | 10 day reversal $\beta cat^{istab}$ **+ FASNi** | | |
|---|---|---|---|---|---|---|---|---|
| **ID** | **Mean** | **SD** | **Mean** | **SD** | **Fold change vs p52 controls** | **Mean** | **SD** | **Fold change vs p52 controls** |
| **10:0** | 1.49 | 0.34 | 0.94 | 0.56 | 0.63 | 1.51 | 0.39 | 1.01 |
| **12:0** | 7.97 | 0.41 | 2.77 | 1.30 ** | 0.35 | 3.78 | 2.11 * | 0.47 |
| **14:0** | 109.88 | 44.22 | 77.61 | 8.75 | 0.71 | 43.36 | 16.50 * | 0.39 |
| **14:1** | 7.85 | 3.10 | 5.02 | 0.72 | 0.64 | 2.30 | 1.01 * | 0.29 |
| **16:0** | 259.27 | 54.82 | 218.76 | 26.19 | 0.84 | 188.09 | 48.22 | 0.73 |
| **16:1 n-7** | 205.96 | 55.28 | 162.35 | 37.76 | 0.79 | 127.42 | 30.34 | 0.62 |
| **18:0** | 128.03 | 31.84 | 86.31 | 21.25 | 0.67 | 97.09 | 33.63 | 0.76 |
| **18:1** | 167.02 | 18.95 | 149.48 | 8.97 | 0.90 | 165.74 | 40.48 | 0.99 |
| **18:2 n-6** | 77.70 | 1.23 | 77.23 | 5.77 | 0.99 | 59.60 | 7.02 ** ## | 0.77 |
| **18:3 n-6** | 1.34 | 0.48 | 1.64 | 0.26 | 1.22 | 1.34 | 0.25 | 1.00 |
| **18:3 n-3** | 18.02 | 4.64 | 22.22 | 2.75 | 1.23 | 16.91 | 6.10 | 0.94 |
| **20:1 n-6** | 35.28 | 12.04 | 28.96 | 8.94 | 0.82 | 23.47 | 6.15 | 0.67 |
| **20:2 n-6** | 7.95 | 2.51 | 6.17 | 1.15 | 0.78 | 7.66 | 2.89 | 0.96 |
| **20:3 n-6** | 1.85 | 0.37 | 1.24 | 0.09 * | 0.67 | 1.17 | 0.21 * | 0.63 |
| **20:4 n-6** | 44.45 | 6.96 | 42.65 | 4.17 | 0.96 | 32.97 | 3.74 * | 0.74 |
| **20:3 n-3** | 6.53 | 1.75 | 5.51 | 0.21 | 0.84 | 6.17 | 1.04 | 0.94 |
| **20:5 n-3** | 4.34 | 0.68 | 3.83 | 0.42 | 0.88 | 1.81 | 0.22 *** ## | 0.42 |
| **22:4 n-6** | 2.82 | 0.54 | 3.46 | 0.67 | 1.23 | 3.32 | 1.08 | 1.18 |
| **22:5 n-6** | 2.36 | 0.52 | 2.90 | 1.02 | 1.23 | 1.56 | 0.14 # | 0.66 |
| **22:5 n-3** | 8.51 | 1.70 | 8.35 | 1.12 | 0.98 | 5.12 | 0.81 * # | 0.60 |
| **22:6 n-3** | 33.47 | 7.43 | 33.71 | 6.84 | 1.01 | 21.76 | 2.93 | 0.65 |
| **MCFA** | 120.45 | 45.83 | 83.18 | 10.50 | 0.69 | 49.15 | 19.54 * | 0.41 |
| **SatFA** | 507.75 | 108.19 | 388.25 | 44.62 | 0.76 | 334.33 | 82.12 * | 0.66 |
| **MUFA** | 408.26 | 68.07 | 340.79 | 39.00 | 0.83 | 316.63 | 66.92 | 0.78 |
| **PUFA** | 209.34 | 24.78 | 208.90 | 7.90 | 1.00 | 159.39 | 17.91 * # | 0.76 |
| **LC-PUFA** | 112.28 | 19.89 | 107.81 | 12.55 | 0.96 | 81.54 | 11.25 * | 0.73 |
| **n-6 PUFA** | 138.46 | 11.25 | 135.28 | 5.33 | 0.98 | 107.62 | 14.37 * # | 0.78 |
| **n-3 PUFA** | 70.88 | 13.83 | 73.62 | 4.11 | 1.04 | 51.77 | 3.97 * # | 0.73 |
| **n6:n3** | 1.98 | 0.23 | 1.84 | 0.10 | 0.93 | 2.07 | 0.16 | 1.05 |
| **AA:EPA+DHA** | 1.18 | 0.04 | 1.15 | 0.12 | 0.97 | 1.41 | 0.22 | 1.20 |
| **SatFA:MUFA** | 1.24 | 0.12 | 1.15 | 0.19 | 0.93 | 1.05 | 0.07 | 0.85 |
| **SatFA:MUFA+PUFA** | 0.82 | 0.09 | 0.71 | 0.10 | 0.87 | 0.70 | 0.07 | 0.85 |
| **18:1/18:0** | 1.35 | 0.33 | 1.80 | 0.44 | 1.33 | 1.74 | 0.18 | 1.29 |
| **16:1/16:0** | 0.79 | 0.06 | 0.74 | 0.13 | 0.94 | 0.68 | 0.03 | 0.86 |
| **20:4n-6/20:3n-6** | 24.16 | 1.27 | 34.45 | 1.92 *** | 1.43 | 28.48 | 2.56 * # | 1.18 |
| **18:3n-6/18:2n-6** | 0.02 | 0.01 | 0.02 | 0.00 | 1.24 | 0.02 | 0.00 | 1.30 |
| **14:1/14:0** | 0.07 | 0.00 | 0.06 | 0.01 | 0.91 | 0.05 | 0.01 | 0.75 |
| **Total FA** | 1125.35 | 195.61 | 937.94 | 52.37 | 0.83 | 810.35 | 164.47 * | 0.72 |

*** Significant with respect to p52 Controls, one-way Anova**

### Significant with respect to 10 day reversal $\beta cat^{istab}$ **+ vehicle, one-way Anova**

**All values in pmoles/mg tissue**

**Table 2.2** Targeted lipidomics analysis on the total fatty acid (esterified + non-esterified) lipid fraction from dorsal skin of p52 controls, vehicle-treated 10-day reversal $\beta$cat$^{istab}$ and FASN inhibited 10-day reversal $\beta$cat$^{istab}$

| | **p52 Controls** | | 10 day reversal $\beta$cat$^{istab}$ **+ vehicle** | | | | 10 day reversal $\beta$cat$^{istab}$ **+ FASNi** | | | |
|---|---|---|---|---|---|---|---|---|---|---|
| **ID** | **Mean** | **SD** | **Mean** | **SD** | | **Fold change vs p52 controls** | **Mean** | **SD** | | **Fold change vs p52 controls** |
| **10:0** | 333 | 236 | 50 | 9 | * | 0.15 | 124 | 53 | | 0.37 |
| **12:0** | 653 | 359 | 159 | 59 | * | 0.24 | 423 | 66 | | 0.65 |
| **14:0** | 2268 | 379 | 1910 | 230 | | 0.84 | 2106 | 282 | | 0.93 |
| **14:1** | 668 | 325 | 467 | 53 | | 0.70 | 466 | 59 | | 0.70 |
| **16:0** | 15105 | 9328 | 12503 | 1759 | | 0.83 | 16184 | 3979 | | 1.07 |
| **16:1 n-7** | 12853 | 6243 | 10511 | 1320 | | 0.82 | 13064 | 2660 | | 1.02 |
| **18:0** | 2358 | 769 | 2217 | 352 | | 0.94 | 2932 | 626 | | 1.24 |
| **18:1** | 4080 | 1552 | 4199 | 530 | | 1.03 | 5426 | 1037 | | 1.33 |
| **18:2 n-6** | 1254 | 302 | 1376 | 290 | | 1.10 | 1342 | 269 | | 1.07 |
| **18:3 n-6** | 101 | 46 | 113 | 24 | | 1.12 | 109 | 34 | | 1.09 |
| **18:3 n-3** | 341 | 14 | 525 | 195 | | 1.54 | 402 | 107 | | 1.18 |
| **20:1 n-6** | 1589 | 758 | 1443 | 493 | | 0.91 | 1346 | 179 | | 0.85 |
| **20:2 n-6** | 328 | 192 | 198 | 74 | | 0.60 | 274 | 66 | | 0.83 |
| **20:3 n-6** | 32 | 20 | 23 | 6 | | 0.70 | 26 | 7 | | 0.81 |
| **20:4 n-6** | 308 | 152 | 355 | 70 | | 1.15 | 340 | 79 | | 1.10 |
| **20:3 n-3** | 113 | 81 | 88 | 28 | | 0.78 | 110 | 24 | | 0.98 |
| **20:5 n-3** | 46 | 30 | 45 | 16 | | 0.99 | 32 | 1 | | 0.69 |
| **22:4 n-6** | 24 | 12 | 32 | 7 | | 1.33 | 37 | 8 | | 1.55 |
| **22:5 n-6** | 24 | 15 | 29 | 10 | | 1.21 | 21 | 3 | | 0.85 |
| **22:5 n-3** | 68 | 39 | 71 | 15 | | 1.03 | 62 | 6 | | 0.90 |
| **22:6 n-3** | 406 | 252 | 355 | 82 | | 0.87 | 276 | 38 | | 0.68 |
| **MCFA** | 3255 | 974 | 2119 | 289 | | 0.65 | 2654 | 363 | | 0.82 |
| **SatFA** | 20718 | 11022 | 16838 | 2242 | | 0.81 | 21769 | 4961 | | 1.05 |
| **MUFA** | 18523 | 8494 | 16153 | 2303 | | 0.87 | 19836 | 3857 | | 1.07 |
| **PUFA** | 3047 | 1148 | 3210 | 787 | | 1.05 | 3033 | 528 | | 1.00 |
| **LC-PUFA** | 1352 | 792 | 1196 | 280 | | 0.89 | 1179 | 160 | | 0.87 |
| **n-6 PUFA** | 2072 | 737 | 2125 | 469 | | 1.03 | 2149 | 430 | | 1.04 |
| **n-3 PUFA** | 976 | 411 | 1085 | 320 | | 1.11 | 883 | 100 | | 0.91 |
| **n6:n3** | 2.16 | 0.13 | 1.99 | 0.16 | | 0.92 | 2.42 | 0.22 | # | 1.12 |
| **AA:EPA+DHA** | 0.71 | 0.08 | 0.89 | 0.03 | | 1.24 | 1.11 | 0.24 | * | 1.55 |
| **SatFA:MUFA** | 1.09 | 0.09 | 1.04 | 0.04 | | 0.96 | 1.09 | 0.06 | | 1.00 |
| **SatFA:MUFA+PUFA** | 0.93 | 0.10 | 0.87 | 0.02 | | 0.93 | 0.95 | 0.05 | | 1.01 |
| **18:1/18:0** | 1.71 | 0.11 | 1.91 | 0.23 | | 1.12 | 1.86 | 0.07 | | 1.09 |
| **16:1/16:0** | 0.91 | 0.15 | 0.84 | 0.03 | | 0.93 | 0.81 | 0.06 | | 0.90 |
| **20:4n-6/20:3n-6** | 10.20 | 2.46 | 15.96 | 1.33 | * | 1.56 | 13.14 | 2.81 | | 1.29 |
| **18:3n-6/18:2n-6** | 0.08 | 0.02 | 0.08 | 0.00 | | 1.06 | 0.08 | 0.01 | | 1.03 |
| **14:1/14:0** | 0.28 | 0.10 | 0.24 | 0.00 | | 0.86 | 0.22 | 0.03 | | 0.79 |
| **Total FA** | 42288 | 20620 | 36201 | 5211 | | 0.86 | 44638 | 9182 | | 1.06 |

*** Significant with respect to p52 Controls, one-way Anova**

### Significant with respect to 10 day reversal $\beta$cat$^{istab}$ **+ vehicle, one-way Anova**

**All values in pmoles/mg tissue**
